# Evidence that viruses, particularly SIV, drove genetic adaptation in natural populations of eastern chimpanzees

**DOI:** 10.1101/582411

**Authors:** Joshua M. Schmidt, Marc de Manuel, Tomas Marques-Bonet, Sergi Castellano, Aida M. Andrés

## Abstract

All four subspecies of chimpanzees are endangered. Differing in their demographic histories and geographical ranges within sub-Saharan Africa, they have likely adapted to different environmental factors. We show that highly differentiated SNPs in eastern chimpanzees are uniquely enriched in genic sites in a way that is expected under recent adaptation. These sites are enriched for genes that differentiate the immune response to infection by simian immunodeficiency virus (SIV) in natural vs. non-natural host species. Conversely, central chimpanzees exhibit selective sweeps at the cytokine receptors *CCR3*, *CCR9* and *CXCR6* – paralogs of *CCR5* and *CXCR4,* the two major receptors utilized by HIV to enter human cells. Thus, we infer that SIV may be eliciting distinctive adaptive responses in different chimpanzee subspecies. Since central chimpanzee SIV is the source of the global HIV/AIDS pandemic, understanding the mechanisms that limit pathogenicity of SIV in chimpanzees can broaden our understanding of HIV infection in humans.

---

Chimpanzees (*Pan troglodytes*) are, alongside bonobos, human’s closest living relatives – the *Pan* and *Homo* lineages having diverged ~6Myr ago (Prado-Martinez et al. 2013). With a genetic divergence of only ~1% (Consortium et al. 2005), *Pan* and *Homo* also share large aspects of their physiology and behaviour, including susceptibility to some pathogens. Studying chimpanzees can teach us about our species by putting recent human evolution in its evolutionary context i.e. the mode and tempo of adaptation and the pressures driving it.

Selection imposed by pathogens has greatly shaped the long-term history of genetic adaptation in the great apes, including chimpanzees and humans (Cagan et al. 2016, Enard et al. 2016). The interest in recent human evolution (Sabeti et al. 2002, Voight et al. 2006, Sabeti et al. 2007, Yi et al. 2010, Racimo 2016) means that we now also have good catalogues of the main targets of local adaptation in many non-African human populations. Nevertheless, analyses of genome-wide patterns of diversity suggest that adaptation via hard selective sweeps has had a limited role in shaping human genomes. Complete selective sweeps involving non-synonymous substitutions appear to have been rare (Hernandez et al. 2011) – but perhaps still important (Enard, Messer, and Petrov 2014). Further, local adaptation has had little effect on the patterns of population differentiation (Coop et al. 2009), unless inferences are boosted with ancient DNA (Key et al. 2016). The focus on humans biases our view on the influence of genetic adaptations in natural populations of primates, and we do not know whether positive selection plays a similarly limited role in shaping other primate genomes. We aim to address this limitation by exploring the recent adaptive history of chimpanzees.

There are four subspecies of chimpanzees, with common names reflecting their location in western and central sub-Saharan Africa: eastern, central, Nigeria-Cameroon and western (Figure 1). Each chimpanzee subspecies is currently endangered, with western chimpanzees critically so (Humle et al. 2016). Subspecies are clearly differentiated, with divergence times ranging from 450 kya to 100 kya, and estimated long-term *N_e_* from 8,000 to 30,000 reflected in varying levels of genetic diversity (Figure 1). There is a wide range of ecological variation across the chimpanzee range, which spans over 5,000 km in sub-Saharan Africa and includes deep forest and savanna-woodland mosaics. Pathogen incidence can also vary between these groups, as seen recently with the lethal outbreaks of Anthrax (Leendertz et al. 2004) and Ebola (Formenty et al. 1999), or the Simian immunodeficiency virus (SIV). SIV, the precursor of the human immunodeficiency virus type 1 (HIV-1) virus that is responsible for the human AIDS pandemic (Keele et al. 2006), is thought to be largely non-lethal to chimpanzees, although some eastern chimpanzees can develop immunodeficiency, see (Rudicell et al. 2010, Keele et al. 2009)). Its prevalence is not uniform across the subspecies, and there is no evidence for infections in western or Nigeria-Cameroon chimpanzees (Locatelli et al. 2016) but multiple infections have been detected in communities of central and eastern chimpanzees (Locatelli et al. 2016, Heuverswyn et al. 2007). Given the separate history and differential environment of each subspecies, and the fact that each subspecies is an independent conservation unit, it is crucial that we identify not only the genetic adaptations shared by all chimpanzees (Cagan et al. 2016), but also the genetic differences conferring differential adaptation to each subspecies.

**Figure 1.**
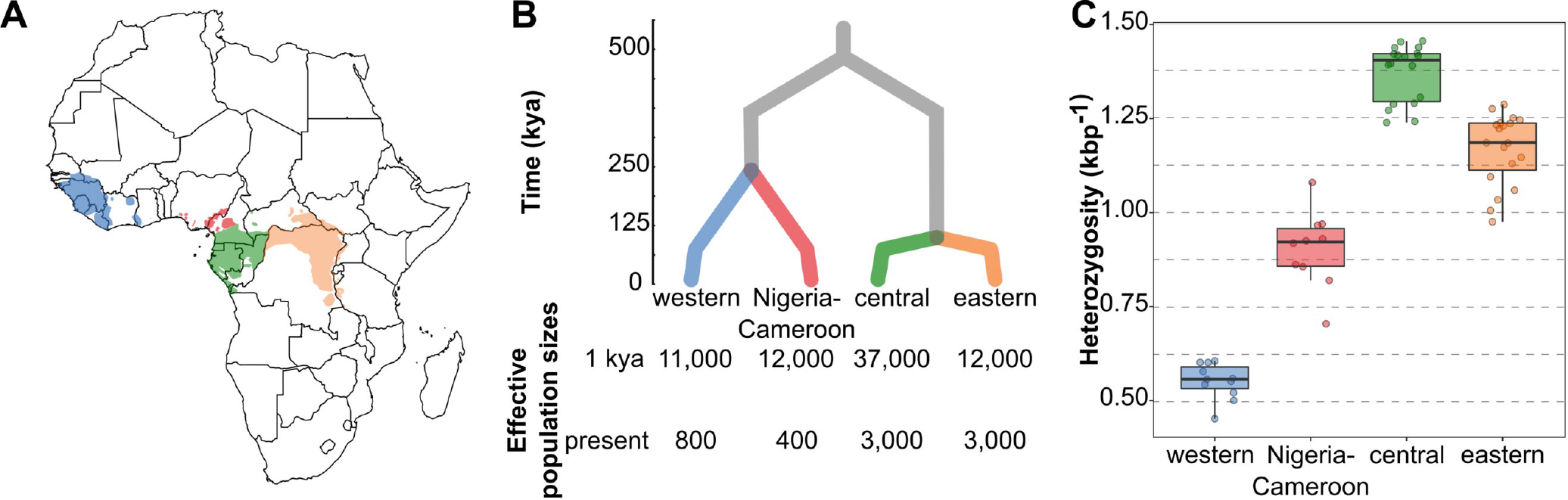
The geographic distribution and population history of chimpanzees. **A**, The ranges of each chimpanzee subspecies within western and central sub-Saharan Africa. Range data from (Humle et al. 2016). **B**, Phylogenetic relationships amongst chimpanzees and the timing of their population divergence, modified from (De Manuel et al. 2016). **C**, Heterozygosity, reflective of relative differences in effective population sizes. Box plots show median central interquartile range, whiskers the upper and lower interquartile range. Points show individual heterozygosity. For all panels, colour designates subspecies: Blue = western, red = Nigeria-Cameroon, green = central, orange = eastern. The heterozygosity counts, and code for plotting panel C are contained in Figure 1–Source Data 1.

To do this, we investigated the signatures of recent genetic adaptation in the genomes of the four subspecies. We show that only eastern chimpanzees have a clear genome-wide signal of recent, local positive selection. This adaptation is potentially due to selection on immunity related genes, with evidence consistent with selection imposed by viruses in general, and SIV in particular. In contrast, putative adaptation to SIV in central chimpanzees seems mediated by adaptation in a suite of cell-entry receptors, results which are suggestive of divergent paths of adaptation to a common pathogen.

## Results

### Genic enrichment in the distribution of derived allele frequency differences

To investigate the influence of recent genetic adaptation in chimpanzee subspecies we compared population differentiation at putatively functional sites (genic sites, defined as +-2kb from protein-coding genes) to differentiation at non-functional sites (here non-genic). Natural selection can only act on functional sites (or affect neutral sites tightly linked to functional sites), so differences between functional and non-functional sites can be ascribed to natural selection. After binning every SNP by its signed difference in derived allele frequency between a pair of subspecies (*δ*), for each bin of *δ* we calculated the genic enrichment, defined as the ratio of genic SNPs vs. all SNPs for each bin of *δ*, normalized by the global genic SNP ratio (Coop et al. 2009, Hernandez et al. 2011, Key et al. 2016). This strategy has been deployed in the study of human local adaptation (Coop et al. 2009, Hernandez et al. 2011, Key et al. 2016), and by not relying on the patterns of linked variation it is not strongly restricted to particular modes of selection. The genic enrichment is greatest for SNPs with the largest *δ*, with the tail bins of *δ* exhibiting significantly greater genic enrichments than any other bin (Figure 2). While not every genic SNP is in this bin due to positive selection, we expect these SNPs, which show the largest frequency differences between subspecies in the genome, to be strongly enriched in targets of positive selection that rose fast in frequency in one of the two subspecies (Coop et al. 2009, Hernandez et al. 2011, Key et al. 2016).

**Figure 2:**
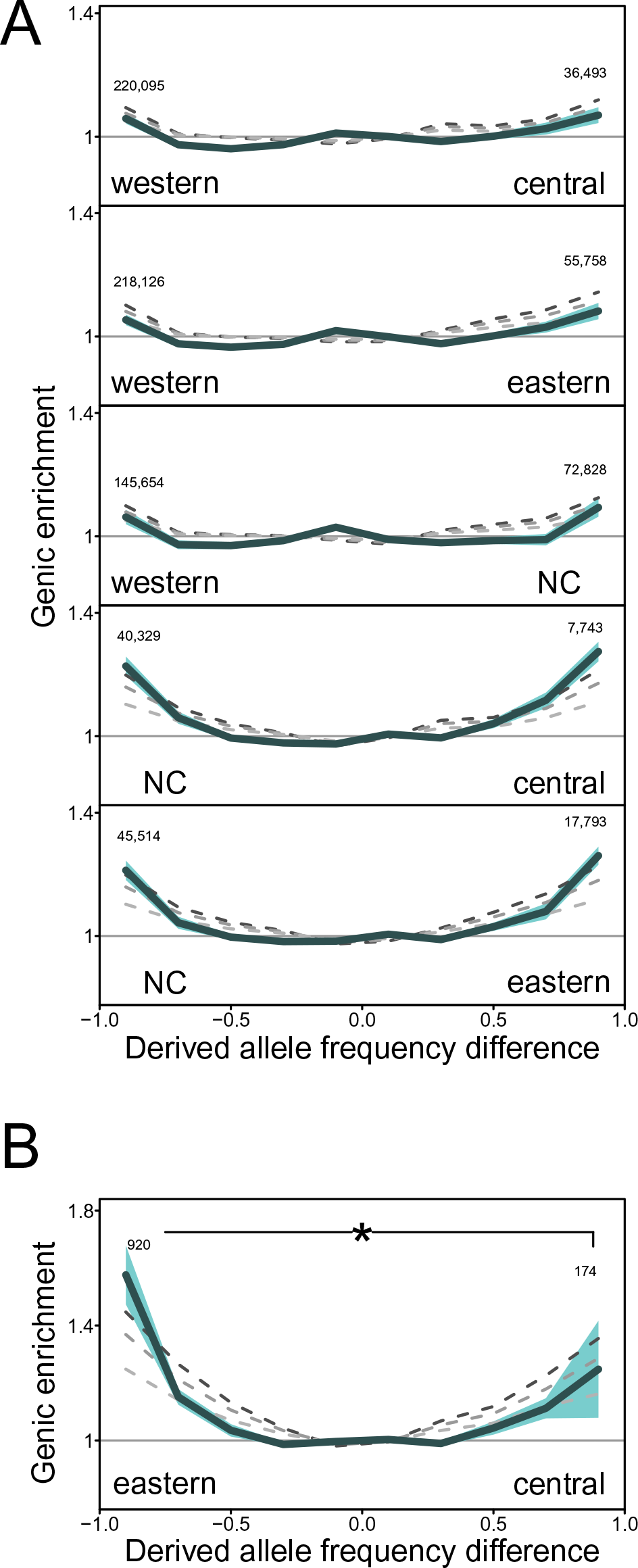
Genic enrichment in bins of signed difference in derived allele frequency (δ). **A,** X-axis: *δ* is computed as the difference in derived allele frequency, for each pair of chimpanzee subspecies. Tail bins (the last bin in either end of *δ*) contain those SNPs with the largest allele frequency differences. Numbers are of the genic SNPs in each tail bin. Y-axis: genic enrichment in each *δ* bin (Methods). **B**, Genic enrichment eastern and central chimpanzee *δ*, plotted separately due to a different Y-axis limit. NC = Nigeria-Cameroon. The asterisk shows significance of the asymmetry in the genic enrichment (* = 0.01). Shading represents the 95% CI (i.e. alpha = 0.05 for a two-tailed test) estimated by 200kb weighted block jackknife. Grey dashed lines represent simulations under increasing levels of background selection that best match different aspects of the data: lightest to darkest shades: B= 0.925 (excluding *δ* tail bins), 0.888 (all *δ* bins), and 0.863 (only *δ* tail bins). The observed and BGS simulated genic enrichments, and code for plotting are contained in Figure 2–Source Data 1.

The genic enrichment in the tails of *δ* is typically roughly symmetric (Figure 2a, symmetry defined as overlapping *δ* tail bin genic enrichment 95% CIs), although the number of tail SNPs and the magnitude of genic enrichment across subspecies pairs varies in accordance with their *N_e_* and divergence times (Figure 2 and Supplementary file 1). Calculated against western chimpanzees, the subspecies with the lowest long-term *N_e_* (de Manuel et al. 2016, Prado-Martinez et al. 2013), the *δ* tail genic enrichment is the least, ranging from 1.05 to 1.09 (Figure 2a). A greater tail genic enrichment, 1.21 to 1.27, is seen for *δ* calculated using Nigeria-Cameroon, the species with the second lowest long-term *N_e_* (Figure 2a). This is comparable to the magnitude of the genic enrichment in the tails of *δ* between human populations (Appendix 1, Appendix 1-figure 1; see (Coop et al. 2009, Hernandez et al. 2011, Key et al. 2016); the genic enrichment across each bin of *δ* also resembles those observed in human populations (Appendix 1, Appendix 1-figure 1; see (Coop et al. 2009, Hernandez et al. 2011, Key et al. 2016).

In marked contrast to these symmetric enrichments, we find a distinctive asymmetry between the tail bin genic enrichments of central and eastern chimpanzees (Figure 2b). The central *δ* tail exhibits a typical genic enrichment (1.25) but surprisingly, the eastern *δ* tail has a genic enrichment (1.58) that is significantly greater than the central tail (0.01 < P < 0.005; weighted 200kb block jackknife, see Methods) and any other *δ* tail (all P < 0.0001; weighted 200kb block jackknife).

The large confidence interval of the central chimpanzee *δ* tail genic enrichment is largely due to the low number but high linkage of SNPs. For example, we found a highly unusual 200kb genomic block on chromosome 3 that contains 70 highly differentiated alleles between central and eastern chimpanzees, similarly distributed among the two tails (37 SNPs in the Central tail and 33 SNPs in the Eastern tail). Concerned that this block could bias our results, we repeated the enrichment analysis after excluding all SNPs contained within it. Removing this block reduces the genic enrichment slightly in the Eastern tail (1.55) but substantially in the Central tail (1.10) resulting in an even stronger asymmetry among the tails. Results are also robust to the removal of the next largest block (with 27 SNPs in the two tails).

To directly quantify asymmetry of the eastern and central chimpanzee *δ* tail genic enrichments, we tested if the log_2_ ratio of each pair of *δ* tail bin genic enrichments departs from zero. Typically, the genic enrichment is greater for the subspecies with the higher long-term *N_e_*. However, the log_2_ ratios are similar and small (ranging from 0 and 0.055), except for eastern vs. central, where it is 0.337 (95% CI, 0.153-0.521, 200kb weighted block jackknife). This is six times larger than the highest ratio between other subspecies pairs (Figure 3; Supplementary file 2, *p-value δ* western vs. central = 0.13 all other *p-values* <= 0.002, *z-test*). The eastern vs. central asymmetry in genic enrichment is thus a clear outlier (*p-value* < 2.2e-16, two-sided Kolmogorov-Smirnov test).

**Figure 3:**
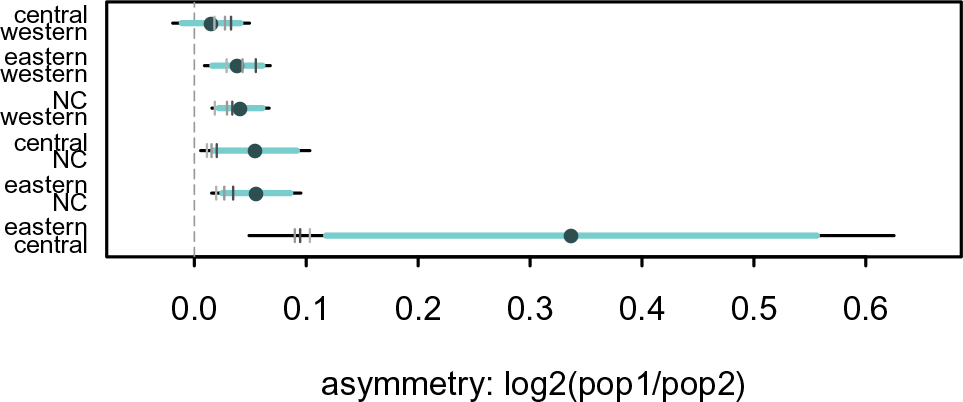
Direct quantification of δ tail bin genic enrichment asymmetry. The asymmetry of the genic enrichments in the *δ* tails is measured by taking their log_2_ ratio, thus 0 indicates a symmetric enrichment (equal enrichment in both *δ* tails). NC = Nigeria-Cameroon. Dot = observed asymmetry. Horizontal lines represent confidence intervals estimated by 200kb weighted block jackknife (light = 95%, black = 99%, i.e. alpha = 0.05 or 0.01 for a two-tailed test). Grey vertical marks represent the *δ* tail asymmetry in simulations, under increasing levels of background selection that best match different aspects of the data: lightest to darkest shades: B= 0.925 (excluding *δ* tail bins), 0.888 (all *δ* bins), and 0.863 (only *δ* tail bins). The observed and BGS simulated genic enrichment tail bin log 2 ratios, and code for plotting are contained in Figure 3–Source Data 1.

### Background selection does not explain the *δ* tail asymmetry

A certain level of *δ* tail bin genic enrichment (Figure 2) is, in principle, compatible with both recent positive selection and background selection (BGS) (Coop et al. 2009), the latter because linkage to sites under purifying selection reduces *N_e_* locally in genic regions and increases the effects of random genetic drift on neutral sites (Charlesworth, Morgan, and Charlesworth 1993). BGS can, for example, explain the *δ* tail bin genic enrichment in human populations, suggesting that this pattern is not evidence for pervasive recent human adaptation (Coop et al. 2009, Hernandez et al. 2011, Key et al. 2016). To explore if BGS can explain our observations, we used coalescent simulations (Methods) to estimate the expected reduction of neutral diversity due to BGS, quantified as a genome-wide average B value (McVicker et al. 2009) that best explains the genic enrichments across all bins of *δ* (the B value that minimizes the summed square differences between observed and simulated enrichments across all pairwise *δ* bins, Appendix 2). This B value is 0.888 – i.e. a reduction of diversity of ~ 11 per cent – decreasing to 0.925 (weaker BGS) when excluding the *δ* tail bins, and increasing to 0.863 (stronger BGS) when fitting solely the twelve *δ* tail bins (Appendix 2, Supplementary file 4). These values agree well with those inferred in humans using similar approaches (Hernandez et al. 2011, Key et al. 2016). It is nonetheless clear that the *B* value of 0.863 that explains the *δ* tail bin genic enrichments results in an extremely poor fit to the genic enrichments in all other *δ* bins (Figure 2).

**Figure 4:**
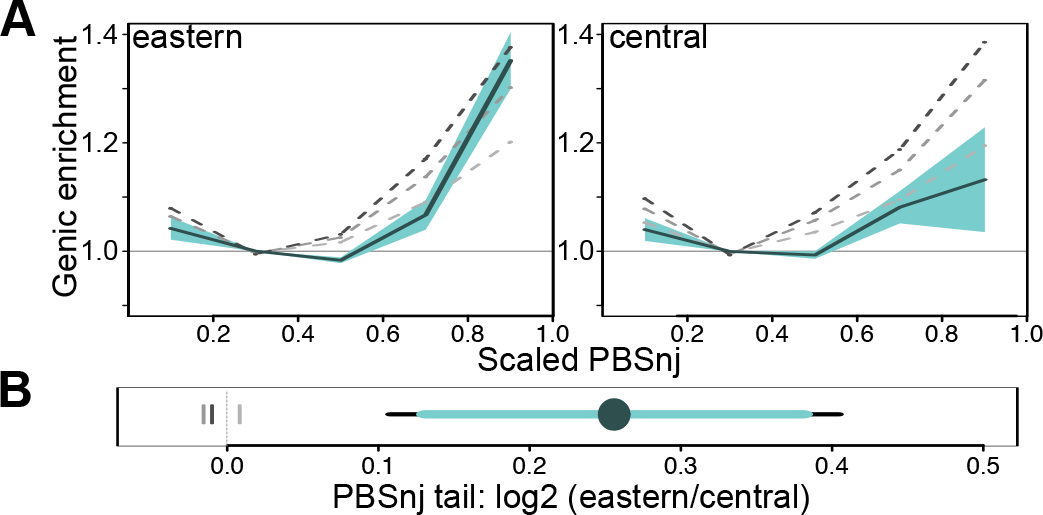
Genic enrichment in bins of PBSnj in eastern and central chimpanzees. **A** X-axes: PBS scaled to take values in the range 0 −1. Y-axes: Genic enrichment computed as described in Figure 2. Shading represents the 95% CI (i.e. alpha = 0.05 for a two-tailed test) estimated by 200kb weighted block jackknife. **B:** log_2_ ratio of the eastern and central PBSnj tail (PBS >= 0.8) genic enrichment. **A,B** Grey dashed (**A**) or vertical (**B**) lines represent the PBSnj genic enrichment in simulations, under increasing levels of background selection that best match different aspects of *δ*, as described in Figs. 2 and 3: lightest to darkest shades: B= 0.925 (excluding *δ* tail bins), 0.888 (all *δ* bins), and 0.863 (only *δ* tail bins). The observed and BGS simulated PBSnj genic enrichments, tail bin log2 ratios, and code for plotting are contained in Figure 4–Source Data 1.

Previously, it was shown that BGS alone does not produce *δ* tail bin genic enrichment asymmetries in comparisons of human populations (Key et al. 2016). We also find that BGS does not result in significant eastern vs. central *δ* tail bin genic enrichment asymmetry. Simulations show a slight asymmetry in the tail genic enrichment (Figs. 2B, 3) due to differences in their demographic histories (Appendix 2, Supplementary file 5). Nevertheless, no simulated value of B in the reasonable range identified above (0.925 – 0.863) results in a tail genic enrichment log_2_ ratio that falls within the 95% CI of the observed ratio (Figure 3). In contrast, the small (though statistically significant) asymmetries in other pairwise *δ* tail bin genic enrichments are observed in simulations and thus fully explicable by demography and BGS (Figure 3). Further, while we find one B value, B = 0.850, that results in a genic enrichment that lies within the 95% CIs for both eastern and central chimpanzees, this B value is a very poor fit to the genic enrichment in all other *δ* bins (Supplementary file 4) and cannot explain the *δ* tail bin genic enrichment asymmetry (observed = 0.337, simulated B of 0.850 = 0.104).

Previous work has shown that background selection varies little among the great apes (Nam et al. 2017). Theory suggests that the diversity reducing effect of BGS is independent of *N_e_*, being determined by the distribution of fitness effects (*s*), except for the narrow range of *N_e_* * *s* = 1 (Nam et al., 2017), while previous work suggests that more than 80% of deleterious mutations in chimpanzees have *N_e_s* » 1 (Bataillon et al. 2015) Thus, the expectation is that the diversity reducing effect of BGS should be the same across all four chimpanzee sub-species. Indeed, we find comparable effects of background selection across subspecies: the relative reduction in neutral variation linked to genes is comparable amongst chimpanzee subspecies (Appendix 3-figure 1a), and neutral diversity has similar dependency on recombination rate and density of functional features across subspecies (with the exception of western chimpanzees, Appendix 3-figure 1b). Further, using a population genetic statistical model (Corbett-Detig, Hartl, and Sackton 2015) we estimate the same reduction in neutral diversity due to background selection in each chimpanzee subspecies, at 11%, in the highest likelihood model (Appendix 3, Supplementary file 6). Thus, despite their differing demographic histories (Figure 1), the effects of BGS are very similar across each chimpanzee subspecies. This justifies using the same average strength of BGS across subspecies above. Nevertheless, to explore if our conclusions are robust to this assumption, we also modelled a greater strength of BGS in eastern chimpanzees (B = 0.825, the value which best matches the eastern *δ* tail bin genic enrichment) than in the other subspecies (B range 0.900-0.850). Stronger BGS in eastern chimpanzees does not produce an eastern central *δ* tail bin asymmetry as large as that observed in the genomes (log2 ratio range 0.120 – 0.146), further illustrating that BGS cannot explain the greater tail genic enrichment in eastern chimpanzees (Figure 3-figure supplement 1). Rather, this is most likely a signal of recent adaptation.

### Population-specific branch lengths with PBSnj

Pairwise comparisons cannot determine which subspecies has changed. Direction, and therefore biological meaning, to allele frequency difference can only be garnered by assuming that derived alleles most often provide the basis for new adaptations. This approach is also limited by the collapsing of the shared history of lineages. For example, in the Nigeria-Cameroon vs. Eastern comparison, 22% of the SNPs in the Eastern *δ* tail are also in the Central *δ* tail (for Nigeria-Cameroon vs. Central comparison), whereas only 3.5% (616 of 17,793) are highly differentiated to both Nigeria and Central chimpanzees. Thus, *δ* summarises the allele frequency change across several parts of the phylogeny, hampering the biological interpretation of its tails.

To overcome this limitation, we developed a statistic that extends the widely used Population Branch Statistic (PBS) (Yi et al. 2010). Briefly, large PBS values identify targets of positive selection as SNPs with population-specific allele frequency differentiation, as these sites result in unusually long branch lengths in pairwise F_ST_-distance trees between three taxa. Small PBS values are due to very short branches, for example due to purifying, shared balancing selection or rare mutations. We extend this test to more than three taxa in the novel *PBSnj* statistic by applying the Neighbor-Joining (NJ) algorithm on the matrix of the per-SNP pairwise F_ST_ distances of the four subspecies (Methods, Appendix 4). This way, PBSnj allows us to jointly compare the four subspecies and identify SNPs with very long branches (allele frequency differentiation) in one subspecies only. Additional advantages of PBSnj are that it does not rely on the specification of ancestral or derived states, and that the NJ algorithm does not require specification of a phylogenetic tree describing the relationship amongst taxa (Appendix 4).

PBSnj allows us to determine within which lineage, eastern or central chimpanzees, allele frequencies have changed to result in the asymmetric *δ* genic enrichment. Analogous to the *δ* tail bins, we binned PBSnj scores and calculated the genic enrichment for each species PBSnj tail (Figure 4A). The PBSnj eastern tail has significantly stronger genic enrichment than the central tail (eastern: 1.36, central: 1.13, log_2_ ratio = 0.25, *p* < 0.001 estimated from weighted 200kb block jackknife, Figure 4B). This shows that the central vs. eastern asymmetry in the *δ* tail bin genic enrichments (Figs. 2B, 3) is due to the drastic allele frequency rise of genic SNPs in eastern chimpanzees since their divergence with central chimpanzees. Importantly, across the range of B values (1.0 – 0.80), simulations show that eastern and central chimpanzee PBSnj tail genic enrichments are expected to be equal (Figure 4B). In fact, BGS would need to be much stronger in eastern chimpanzees than in central chimpanzees to produce the observed levels of PBSnj tail genic enrichments. BGS with B < 0.888 would be required to produce the genic enrichment exhibited in the eastern PBSnj tail, but B = 0.888 produces PBSnj tail genic enrichments of equal or greater magnitude as those seen for central chimpanzees (and also Nigeria-Cameroon and western, Appendix 5 and Appendix 6, and Supplementary file 9). Thus, it is eastern chimpanzees that exhibit the greatest genic enrichment for highly differentiated SNPs, an enrichment that (unlike in other subspecies) we cannot explain by background selection alone. This suggests the greater enrichment in the PBSnj eastern tail is due to positive selection, and by using the genomic blocks used to estimate the PBSnj tail Confidence Intervals in Figure 4A, we estimate that an additional eight-19 population specific sweeps are sufficient to explain this difference (Methods, Figure 4-figure supplement 1). Although this is a conservative estimate, it shows that we do not require an unrealistically large number of selective sweeps to explain the distinct pattern of eastern chimpanzees.

### Long-range LD and regulatory functions in the PBSnj eastern tail SNPs

Further evidence that the PBSnj eastern tail genic enrichment is not due to background selection would be provided by independent signatures such as the patterns of linkage disequilibrium (LD) and the putative functional consequences of alleles. LD based tests of positive selection are more robust to background selection than those based on population differentiation (Enard, Messer, and Petrov 2014). We computed three haplotype-based selection statistics that identify the signatures of positive selection within populations (*i*HS; (Voight et al. 2006), nSl (Ferrer-Admetlla et al. 2014)) or between populations (*XP-EHH* (Sabeti et al. 2007)). For each statistic, PBSnj eastern tail SNPs have a significantly higher score than randomly sampled genic SNPs (mean *i*HS 0.69, mean nSl 0.94, and mean XP-EHH mean 0.51, standardized for the genic background to have mean = 0 and sd = 1 for each statistic; all *p* < 0.0001; re-sampling test; Supplementary file 10). Thus, SNPs specifically differentiated in the eastern PBSnj tail have on average higher LD-based signatures of recent positive selection than random genic SNPs, and also to a greater degree than all other subspecies’ PBSnj tails (Supplementary file 10) (two-sample *t-*tests, all *p-values* << 0.0001).

PBSnj tail genic SNPs are significantly enriched in exonic variants, but not in non-synonymous (as compared with synonymous) ones, so less than 1% of PBSnj eastern tail SNPs result in amino acid changes (observed = 0.84%; genic background = 0.18 %, p < 0.001, Supplementary file 11 lists PBSnj eastern tail non-synonymous SNPs). Turning to regulatory changes, we used regulomeDB (Boyle et al. 2012) to predict putative regulatory consequences of chimpanzee SNPs from the sequence context and biochemical signatures of homologous human sites. The PBSnj eastern tail genic SNPs are more likely to have strong evidence of regulatory function (3.7% vs. 3.0%, permutation test p = 0.012) and less likely to have no ascribed regulatory function (52.3% vs. 56.0%, permutation test p = 0.0001) than randomly sampled genic SNPs, Supplementary file 12. In contrast, PBSnj central tail SNPs show no difference to the genic background for either category (Supplementary file 12; Nigeria-Cameroon and western also exhibit weaker but significant enrichments). Interestingly, PBSnj eastern tail SNPs do not differ in functional constraint (as measured by *phastCons* scores (Siepel et al. 2005, see Methods) from random genic SNPs (Supplementary file 13). This suggests that while likely enriched in regulatory functions, these sites are not under particularly strong long-term constraint, perhaps because they do not affect functions that have been tightly conserved over long evolutionary times.

### Biological functions of the PBSnj eastern tail SNPs

To understand the biological mechanisms and putative selective factors driving the recent adaptations in eastern chimpanzees, we investigated the genes containing the genic SNPs in the PBSnj eastern tail (hereafter PBSnj eastern genes). Two Gene Ontology (GO) categories (Ashburner et al. 2000, The Gene Ontology 2017) are significantly enriched (p < 0.05, False Discovery Rate (FDR) < 0.05; GOWINDA; Supplementary files 15-18). The top category is “cytoplasmic mRNA processing body assembly”, and three of the five PBSnj eastern genes in this category (*DDX6* (Ayache et al. 2015), *ATXN2* (Nonhoff et al. 2007) and *DYNC1H1* (Loschi et al. 2009)) are either key components of processing bodies (P-bodies) or regulate the assembly or growth of P-bodies in response to stress. Selection on the immune system is suggested by the second category, “antigen processing and presentation of peptide antigen via MHC class I”. The signal in this category is due to six genes, of which only *HLA-A* is an MHC gene, with the other genes being *B2M*, *ERAP1*, *PDI3*, *SEC13,* and *SEC24B*. With FDR < 0.1, there are three more significant categories related with immunity: “T cell co-stimulation”, “negative regulation of complement-dependent cytotoxicity”, and “type I interferon signalling pathway”. There is thus a preponderance of immunity-related GO categories and genes involved in anti-viral activity (see Discussion). Even the “cytoplasmic mRNA processing body assembly” category is potentially linked to virus infection as P-bodies are cytoplasmic RNA granules manipulated by viruses to promote viral survival and achieve infection (Tsai and Lloyd 2014, Lloyd 2013). Interestingly, the PBSnj eastern genes are also enriched in three sets of Viral Interacting Proteins (VIPs) (see Supplementary files 19-22), which are genes with no annotated immune functions but that interact with viruses (Enard et al. 2016). As VIP sets do not in the main contain classic or known immunity genes, this provides an independent signal for the relevance of viruses in this gene set. Together, these results suggest that adaptation to pathogens, and viruses in particular, may have had an important role in the recent adaptation in eastern chimpanzees.

Amongst chimpanzee viruses, the simian immunodeficiency virus (SIV) is intensively studied as it is the progenitor of the human immunodeficiency virus (HIV) that created the global acquired immune deficiency syndrome (AIDS) pandemic. It is also of interest here because it appears to only infect natural populations of eastern and central chimpanzees (Santiago et al. 2002, Santiago et al. 2003, Nerrienet et al. 2005, Boué et al. 2015), and because it has mediated fast, recent adaptations in other natural hosts (Svardal et al. 2017). Svardal *et. al.* (2017) investigated a set of genes that change expression in response to SIV infection in SIV natural hosts (vervet monkeys) but not in non-natural hosts that develop immunodeficiency (macaques) (Jacquelin et al. 2014, Jacquelin et al. 2009), hereafter referred to as “natural host SIV responsive genes”. Natural host SIV responsive genes are likely involved in the specific, early immune response of natural hosts to SIV infection, which limits the effects of the virus and prevents subsequent immunodeficiency. These genes show signatures of positive selection in vervet monkeys, suggesting that ongoing adaptation to the virus in natural hosts can occur (Svardal et al. 2017). Strikingly, the PBSnj eastern tail SNPs are significantly enriched in these same natural host SIV responsive genes (Jacquelin et al. 2014, Jacquelin et al. 2009) (observed 118 genes, expected 100, *p-value =* 0.0195, GOWINDA, FDR < 0.1 see Methods, Table 1, Supplementary file 23). In fact, the set of natural host SIV responsive genes can fully explain the unique eastern signature: the asymmetry in the PBSnj tail is abolished when this set of genes is removed from the analysis (genic enrichment in the eastern PBSnj tail decreases from 1.36 to 1.26, and the 95% confidence interval of this point estimate now overlaps those of Nigeria-Cameroon and central chimpanzees (Methods). A reduction in the genic enrichment in the PBSnj tail is expected, as it is enriched in natural host SIV responsive genes; but this exercise allows us to show that in the absence of selection in natural host SIV responsive genes, the signature of recent positive selection in eastern chimpanzees would not be exceptional. The natural host response in vervet monkeys is associated with changes in the expression of these natural host SIV responsive genes. In agreement with potential adaptations in gene expression, the set of PBSnjE SNPs in the natural host SIV responsive genes is further enriched in sites with a high likelihood having an inferred gene regulatory function (p=0.0485 when compared with other PBSnj eastern tail genic SNPs, p=0.0089 with all genic SNPs) and strongly depleted of sites with no predicted regulatory function (p=0.0001 when compared with other PBSnj eastern tail genic SNPs, p=0.0001 with all genic SNPs, Supplementary file 24). While these genes were not identified in chimpanzees, this suggests a similar mechanism of adaptation to SIV (or to an unknown virus with a similar effect in gene expression) in vervet monkeys and chimpanzees.

**Table 1:** VIP gene enrichment in the PBSnj eastern tail.

| **VIRUS** | **P-VALUE** | **FDR P-VALUE** |
| --- | --- | --- |
| BLV | 0.0015 | 0.0239 |
| DENV | 0.0025 | 0.0239 |
| HTLV | 0.0145 | 0.0780 |

**Table 2:** SIV responsive gene enrichment in subspecies PBSnj tails.

| **Subspecies** | **Observed** | **Expected** | **P-VALUE** |
| --- | --- | --- | --- |
| Eastern | 118 | 99 | 0.0198 |
| Central | 36 | 29 | 0.0739 |

### Biological functions of the PBSnj central tail SNPs

Despite having a larger long-term *N_e_* than eastern chimpanzees, central chimpanzees do not show a clear genomic signature of recent adaptation. Despite being naturally infected by SIV and being the source of pandemic HIV, they show no clear indication of selection in SIV responsive genes: the PBSnj central tail has a greater number of SNPs in SIV responsive genes than expected (36 vs. 29), but the enrichment is non-significant (p = 0.0756; resampling test, Table 1). Power to identify a significant enrichment may be hampered by the low number of SNPs. However, highly differentiated SNPs in the PBSnj long branches of central chimpanzees are significantly enriched in one GO category, “chemokine receptor activity”, due to SNPs in *CCR3*, *CCR9* and *CXCR6* (*p =* 0.00001, FDR = 0.0197, GOWINDA). Each of these genes is located within the large cluster of cytokine receptor genes on chromosome 3, but they appear to be associated with different sweep events (Figure 4-supplement 3). These genes are of interest because *CCR3* and *CXCR6* have paralogs (*CCR5* and *CXCR4*) that in humans are the two most common co-receptors for HIV-1 cell entry (Berger 1997, Moore et al. 2004). Both *CCR3* and *CXCR6* can be used to enter the cell by some SIV, HIV-1 and HIV-2 subtypes (Nedellec et al. 2010, Gorry et al. 2007, Bron et al. 1997, Willey et al. 2003), and the SIV of both Sooty mangabey (Elliott et al. 2015) and Vervet monkey (Wetzel et al. 2017) use *CXCR6*. The breadth of co-receptors used by SIV in chimpanzees is unknown, but sequence changes in the V3 section of the virus can modify the specificity of the co-receptors used by HIV (Gorry et al. 2007). We note that one of the PBSnj tail SNPs in *CCR3* results in an amino acid substitution (246 S/A) in transmembrane domain 6, and this region has been implicated in the modulation of CCR5 activity (Steen et al. 2013). Thus, changes in these co-receptors may have the potential to affect the entry of SIV in chimpanzee cells.

## Discussion

Comparing whole genomes from the four subspecies of chimpanzees we find that the alleles whose frequency rose quickly and substantially in particular chimpanzee subspecies, resulting in strong genetic differentiation, are enriched in genic sites. Except for eastern chimpanzees, these genic enrichments can be explained by the effects of BGS as is the case in human populations (Coop et al. 2009, Hernandez et al. 2011, Key et al. 2016). Our PBSnj statistic shows that this signature in eastern chimpanzees is due to SNPs whose frequency have changed specifically in eastern chimpanzees since their divergence with central chimpanzees. These sites tend to have high long-range LD, but few of them have significant LD signatures of positive selection. Many of them are polymorphic in central chimpanzees, so it is likely that many of these adaptations have occurred from standing genetic variation and consist of soft sweeps (Hermisson and Pennings 2017). This would suggest that adaptation from standing genetic variation is important throughout primate evolution, not just in recent human evolution (Pritchard, Pickrell, and Coop 2010). Alternatively, some of these sites may be polymorphic in central chimpanzees due to gene flow from eastern chimpanzees. The inferred chimpanzee demography includes recurrent migration between eastern and central chimpanzees, in both directions (de Manuel et al. 2016), requiring selection in eastern chimpanzees to be strong enough to overcome the homogenising effect of gene flow.

These strongly differentiated alleles in eastern chimpanzees are enriched in sites with inferred regulatory function, but not in sites that have been strongly constrained during mammalian evolution. This agrees well with a role in adaptation to pathogens, which is often characterized by fast arms-race evolution. The PBSnj eastern genes are enriched in several immune-related categories, with many of them having known or potential virus-related functions. *OAS2* and *RNASEL*, for example, are involved in foreign RNA degradation (Sadler and Williams 2008), while *ERAP1* is a gene under long term balancing selection in humans (Andrés et al. 2010) that is involved in MHC class I epitope presentation (Hearn, York, and Rock 2009). These are plausible adaptations to viral infections in eastern chimpanzees. In fact, these PBSnj eastern sites are located disproportionately in genes that differentiate the CD4 transcriptional response to SIV in a natural host species that tolerates the virus from a non-natural host species that develops immunodeficiency. Selection acting on this set of genes is sufficient to produce the greater eastern signal. Two aspects of this enrichment are notable. First, these genes are identified based on gene expression responses in vervet monkeys and macaques to SIV infection (Jacquelin et al. 2014, Jacquelin et al. 2009), and are thus completely independent of chimpanzee genetics. Second, the SIV responsive genes also show diversifying selection in vervet monkeys (Svardal et al. 2017). Of note, these SNPs are strongly enriched in putative regulatory functions, in agreement with putative adaptations through gene expression. This suggests that SIV may continue to exert an important selective force in natural primate species, which both vervet monkeys and eastern chimpanzees may respond to by shaping gene expression.

How may this happen? The genes that are both SIV-responsive and contain PBSnj eastern tail SNPs are significantly enriched in four GO categories (FDR < 0.1, GOWINDA, Supplementary file 25). The top category is “type I interferon signalling pathway with four genes (*IRF2*, *RNASEL*, *HLA-A* and *SP100*). This category is also significantly enriched in the full set of PBSnj eastern tail SNPs. *OAS2* is also in this category but it is inducible in both vervet and macaque shortly after SIV infection. *IRF2*, *RNASEL* and *SP100* are all upregulated in the CD4 cells of vervet monkey but not of macaques one day post infection. This is relevant as regulation of the interferon response is a key differentiator between natural and non-natural SIV hosts (Harris et al. 2010) and the timing of interferon responses can be key in the progression to AIDS in humans infected with HIV (Rotger et al. 2011, Utay and Douek 2016). Another enriched category is “polycomb group (PcG) protein complex”. PcG complexes can be involved in the epigenetic regulation of HIV-1 latency (Friedman et al. 2011, Khan et al. 2018), and three of the genes in this GO category, *PHC2*, *CBX7* and *KDM2B* encode components of the same PcG complex, PCR1 (Khan et al. 2018). While in vervet monkeys *CBX7* and *PHC2* are both downregulated in CD4 cells six days post infection, *KDM2B* is upregulated 115 days post infection, which might hint at increased epigenetic control of SIV in the chronic phase of infection.

Of course, it is also possible that other viruses elicited a selection response in eastern chimpanzees, and in particular the SIV signature that we observe could be due to selection by other ssRNA viruses. Possibilities include the viruses involved in the three significant sets of VIPs, which are *Dengue virus* and the closely related *Bovine leukaemia virus* and *human T-lymphotropic virus.* Nonetheless, SIV is a better candidate to explain our observations. The set of genes that explains the PBSnj eastern signature, the natural host SIV responsive genes, have a clear functional, direct involvement in response to SIV virus in African primates (Svardal et al. 2017). These genes show signatures of recent positive selection also in vervet monkeys (Svardal et al. 2017), suggesting that SIV is an important selective force even in natural hosts. Chimpanzees have been classically described as natural SIV hosts, but some reports suggest fitness consequences in populations of eastern chimpanzees infected with the virus (Keele et al. 2009), with some infected individuals described as having an AIDS-like pathology. It thus seems likely that the virus is a selective force in this subspecies. Thus, while we cannot be completely certain that SIV is driving selection in eastern chimpanzees, this virus is the best candidate considering all currently available evidence.

It is also probable that eastern chimpanzees have adapted to additional selective pressures unrelated of viral pathogens or immunity. An obvious candidate would be life history traits. For example, the gene *SKOR2*, which contains the fifth ranked eastern specific missense polymorphism, has been associated with the timing of female puberty in human GWAS of age of menarche (Pickrell et al. 2016). Unfortunately, the genetic basis of these traits is poorly understood making it hard to contextualise this result.

Perhaps surprisingly, central chimpanzees have weaker signatures of natural selection despite being the subspecies with the largest *N_e_* (de Manuel et al. 2016, Prado-Martinez et al. 2013). A few factors could blunt the evidence for positive selection in central chimpanzees, but none of them are able to explain the observed difference in PBSnj tail genic enrichment between central and eastern chimpanzees –including putative population substructure, gene flow from eastern chimpanzees or introgression from bonobos (Appendix 7). Central chimpanzees also do not have significant enrichment in SIV responsive genes despite, like eastern chimpanzees, being naturally infected by SIV (Heuverswyn et al. 2007). However, central chimpanzees exhibit a significant enrichment of highly differentiated SNPs in the chemokine genes *CCR3*, *CCR9* and *CXCR6*, paralogs of which are involved with HIV cell entry in humans. *CCR3* and *CXCR6* are used by SIV, HIV-1 and HIV-2 subtypes (Nedellec et al. 2010, Gorry et al. 2007, Bron et al. 1997, Willey et al. 2003, Wetzel et al. 2017, Elliott et al. 2015) and a SNP in *CCR3* may modulate the activity of the channel. The signature of positive selection in *CXCR6* is interesting because the SIV of natural hosts Sooty Mangabey (Elliott et al. 2015) and Vervet monkeys (Wetzel et al. 2017) predominantly use *CXCR6* for host cell entry. This is in contrast with the dominant *CCR5* usage in hosts such as humans and macaques that progress to AIDS. While it is unclear which particular channels are used by SIV in each chimpanzee subspecies, the evidence of selection in central chimpanzees in these receptors raises the intriguing possibility that the two chimpanzee hosts have used strikingly distinct evolutionary responses to the virus: limiting cell entry in central chimpanzees; modulation of gene expression response in eastern chimpanzees. With the estimated time of infection being ~100,000 years ago, this could be due to differential adaptation to a common selective pressure, or potential subspecies-specific coadaptation between chimpanzee hosts and SIV. This last is an intriguing possibility that warrants further investigation.

While our attention has focussed on eastern, and to a lesser extent, central chimpanzees, this is not to say that positive selection has not acted on western and Nigeria-Cameroon chimpanzees. Rather, what we conclude is that, as in the case of central chimpanzees, BGS under the inferred demography of chimpanzees is adequate to explain the patterns of genetic differentiation in these subspecies. We note however that the divergence time of these lineages makes tests of allele frequency differentiation less well suited to identify adaptive loci than in eastern and central chimpanzees. Alternative approaches, for example using intensive within subspecies sampling, can help identify adaptive loci in these subspecies. Nevertheless, our results show striking differences between the sister subspecies of eastern and central chimpanzees. Besides helping us start to identify the genetic and phenotypic differences among subspecies, this finding highlights the need for genetic studies and conservation efforts to account for functional differentiation between subspecies and local populations across the entire chimpanzee range.

## Materials and Methods

### Genotypes, haplotypes and genic regions

We analysed the 58 chimpanzee genomes described in de Manuel *et. al.* (2016), with sample sizes of: eastern 19, central 18, Nigeria-Cameroon 10, western 11 after excluding the hybrid Donald. For most tests based on allele frequencies, we used the chimpanzee VCF file from de Manuel et al., (2016) after removing every SNP with at least one missing genotype across all chimpanzees. For haplotype phasing, we also included the 10 bonobo genomes from (de Manuel et al. 2016). To statistically phase haplotypes we used *Beagle (Browning and Browning 2007) v 4.1* (downloaded from https://faculty.washington.edu/browning/beagle/b4_1.html, May 2016). We used default parameters without imputation, except that after the initial 10 burn in iterations we performed 15 phasing iterations (default is five) using the following command line: java -Xmx12000m -jar beagle.03May16.862.jar gt=vcf out=vcf.phased impute=false nthreads=1 niterations=15

For the analysis of *δ* we chose to use the homologous human genome reference allele as the ancestral state for chimpanzee SNPs. We used the human genome from the 1000 Genomes project phase III human_g1k_v37.fasta, available from: http://ftp.1000genomes.ebi.ac.uk/vol1/ftp/technical/reference/human_g1k_v37.fasta.gz

We used the UCSC liftover utility to convert chimpanzee SNPs’ coordinates from pantro 2.1.4 to human genome version 37 (hg19) coordinates, then used samtools faidx to retrieve the human allele for that position.

We acknowledge that some AA inferences can be incorrect due to parallel mutations in the human lineage or lineage sorting effects. To show that our result was robust to AA inference method, we also used the homologous gorilla allele and parsimony of both the human and gorilla allele extracted from 20 mammalian multiz alignment to the human genome hg38, downloaded from UCSC (http://hgdownload.cse.ucsc.edu/goldenPath/hg38/multiz20way/maf/), and the inferred chimpanzee ancestral allele determined calculated using the EPO alignments and downloaded from ENSMBL, available at ftp://ftp.ensembl.org/pub/release-90/fasta/ancestral_alleles/pan_troglodytes_ancestor_CHIMP2.1.4_e86.tar.gz. Each of these inference methods recovered the same signal of a significantly greater *δ* tail bin genic enrichment for eastern vs. central chimpanzees, see Supplementary file 14. Again, we also note that our new statistic PBSnj does not require inference of the ancestral allele.

We considered protein-coding genes on the autosomes (17,530 genes) and define ‘genic sites’ by extending the transcription start and end coordinates from ENSEMBL biobank for pantro2.1.4 by 2kb on each side.

### Genetic map

For statistics that required a genetic map, we used the pan diversity genetic map (Auton et al. 2012) inferred from 10 western chimpanzees. We downloaded the chimp_Dec8_Haplotypes_Mar1_chr-cleaned.txt files from birch.well.ox.ac.uk/panMap/haplotypes/genetic_map. These files consist of SNPs and their inferred local recombination rate. These map data were inferred from sequences aligned to the pantro2.1.2 genome, so we used two successive liftover steps to convert the coordinates of sites used to infer the genetic map to pantro2.1.4 coordinates: pantro2.1.2 to pantro2.1.3, then pantro2.1.3 to pantro2.1.4. Two steps are required as there are no liftover chains relating pantro2.1.2 to pantro2.1.4. Of the 5,323,278 autosomal markers, 33,263 were not lifted from pantro2.1.2 to pantro2.1.3. The remaining 5,290,015 were also successfully converted to pantro2.1.4 coordinates. After liftover we filtered sites that after the two steps were mapped to unassigned scaffolds or the X chromosome, which left 5,289,844 SNPs. Next, we sorted loci by position to correct cases where their relative order was scrambled. This left a final number of 5,289,460 autosomal SNPs. Recombination rates were then recalculated by linear interpolation between consecutive markers (marker x, marker y) using the average of their estimated recombination rates (rate x, rate y). These recombination maps have been deposited on Dryad (see Data availability).

### Signed difference in derived allele frequency (*δ*)

Using the derived allele frequency of each SNP for each subspecies we calculated, for each pair of chimpanzee subspecies, the signed difference in derived allele frequency (DAF) between them: *δ =* DAF_pop1_ – DAF_pop2_; DAF_pop1_ > DAF_pop2_: *δ* > 0; DAF_pop1_ < DAF_pop2_: *δ* < 0; −1 <= *δ* <= 1. We bin *δ* into 10 bins of 0.2. The choice of subspecies assigned to pop1 or pop2 is arbitrary and has no bearing on the results. To ensure that both tail bins are identically wide, we define them as Bin 1: −1 <= *δ* <= 0.8 and Bin 10 as 0.79 < *δ* <= 1. As a consequence, the Bin 5 (0.00 < *δ* < 0.2) is marginally narrower than the other bins (by 0.01), but it contains a large number of sites and the slight size difference has negligible impact on the analyses. The derived allele counts have been deposited on Dryad (see Data availability).

We estimate confidence intervals and infer *p-values* for *δ* genic enrichment using a weighted block jackknife (Reich et al. 2009) utilising the method of Busing *et. al.* (Busing et al. 1999). This has been used for analogous tests, as it accounts for linkage disequilibrium, which means that SNPs in *δ* bins are not full independent of each other. We divide the genome into non-overlapping 200kb windows to capture the blocking effect of LD. We then recalculate, for each bin, the genic enrichment using a delete-1 window jackknife. We also weight the windows by the total number of SNPs in them, to downweigh, within each bin, blocks with large numbers of linked SNPs. We determine that two tails are differentially enriched if their 95% CIs of enrichment do not overlap. For directly testing asymmetry (or in the case of PBSnj, equality) using the log_2_ ratio, we use the same weighted block jackknife, and use the 95% CI as a two-tailed test with alpha = 0.05. Other enrichment and resampling tests are described in Methods subsection “Statistics”.

### Population Branch Statistic neighbour-joining

The Population Branch Statistic (PBS, Yi et al. 2010) is a test of population specific natural selection. In the framework of a three-taxon distance tree, SNPs under selection specific to one population are detected as those that result in longer than expected branch lengths (large allele frequency differentiation). To generate the tree, for each site, the full distance matrix of pairwise F_ST_ is computed. A three taxa tree is unrooted and has only one possible topology, so simple algebra allows the calculation of each branch length in the tree. Extreme outliers in the distribution of PBS are considered candidates of positive selection.

We introduce Population Branch Statistic neighbour-joining (PBSnj) as a simple method to calculate population specific branch lengths when more than three taxa are being analysed. We note that related Methods have recently appeared in the literature (Cheng, Xu, and DeGiorgio 2017, Racimo, Berg, and Pickrell 2018). Full details are in Appendix 4, but in brief, using the full matrix of pairwise F_ST_, F_ST_ values are transformed to units of drift time as ln (1-F_ST_) (Yi et al. 2010). For fixed differences this transformation is mathematically undefined i.e. ln (0), and F_ST_ =1 is replaced with the next largest observed F_ST_ value for a given population pair. Then the Neighbor-Joining algorithm (Saitou and Nei 1987) is used to infer the tree topology and calculate branch lengths. This overcomes errors in the inferred length of external branches due to misspecification of a fixed tree topology. To enable a binning scheme of PBSnj values that is comparable between subspecies, these scores are further normalised to be on the 0-1 scale. These data have been deposited on Dryad (see Data availability). F_ST_ for PBSnj was calculated using the estimator described in (Bhatia et al. 2013) because there are unequal sample sizes for the subspecies, and the classical Weir and Cockerham estimator can be biased with unequal sample sizes (Bhatia et al. 2013). To calculate genic enrichments along the PBSnj distribution we bin SNPs in PBSnj bins 0.2 units wide. As for *δ* analyses, we use the 200 kb weighted block jack-knife to estimate confidence and significance levels. We provide a source code file, written in R, to calculate PBSnj (“PBSnj_method.R”).

### Model of Chimpanzee demographic history

The most detailed exploration of chimpanzee demography comes from the work of de Manuel *et. al.* (2016). This paper describes the 58 chimpanzee full genome sequences we use here, and estimation of their inferred demographic model. As this paper took a primary interest in investigating chimpanzee-bonobo post speciation gene flow, and to reduce the number of parameters to be estimated, models were inferred using either Nigeria-Cameroon or western chimpanzees, but not both. Thus, de Manuel *et. al.* (2016) provides “bonobo, eastern, central, Nigeria-Cameroon” and “bonobo, eastern, central, western” models. These are referred to, respectively, as ‘becn’ and ‘becw’ models below.

For this investigation we use a merged demographic history. To begin the construction of this model, we recognised that there is little gene flow involving western chimpanzees in the ‘becw’ model, but that gene flow events are a key determinant of patterns of chimpanzee genetic diversity and differentiation in the ‘becn’ model. We therefore used the ‘becn’ model as a scaffold to which parameters relating to western chimpanzees (bottlenecks, expansions and *N_e_* estimates) from the ‘becw’ model are “grafted” in, to create a merged ‘becnw’ model. To make sure that the *N_e_* of western chimpanzees was appropriately scaled, all *N_e_*s 1000 ya pastwards for western chimpanzees specified in the ‘becw’ model were normalized by multiplying by the ratio of the inferred *N_e_*s of central chimpanzees specified from 1000 ya pastwards in the ‘becn’ and ‘becw’ models: scaled western *N_e_* = western *N_e_* * 3.66914400056 / 4.3158739382. Present western *N_e_* was normalised by the ratio of the present central *N_e_*: scaled western *N_e_* = western *N_e_* * 0.3092 / 0.30865.

Initially, we used the split time of the western and Nigeria-Cameroon lineages of 250ky reported by de Manuel *et. al*. which was estimated from sequence divergence data, but this gave a bad fit to F_ST_ values, being substantially lower than observed (Supplementary file 3). We addressed this by increasing the western/Nigeria-Cameroon divergence time in proportion to the ratio of model:observed western/Nigeria-Cameroon F_ST_. i.e. F_ST_Observed / F_ST_Model = timeX / 250kya => timeX = F_ST_Observed / F_ST_Model x 250kya. We adjust the observed F_ST_ by −0.008 – to capture the average difference between model versus observed F_ST_ values for central/eastern/Nigeria-Cameroon chimpanzees. This simple calculation results in an adjusted time of 267kya for the western/Nigeria-Cameroon split. F_ST_ values for this new model show a much better fit to observed values (Supplementary file 3), and it is this model that we use for all subsequent modelling of genic enrichments and the effects of background selection.

To determine model fit above, we calculated all pairwise average F_ST_ values for the simulated data and compared them to the empirical F_ST_ estimates. For each scenario, we simulated 1,000,000 2kb fragments (2 Gb of sequence).

All simulations of neutral diversity and background selection were performed with *msms* (Ewing and Hermisson 2010), and following de Manuel *et. al.* assuming a mutation rate of 1.2e^-8^ and recombination rate 0.96e^-8^, with the following command line:

msms 116 1 -t 0.96048 -r 0.768384 2001 -I 5 0 38 36 20 22 0 -n 1 0.0742 -n 2 0.3181 -n 3 0.3092 -n 4 0.0386 -n 5 0.08114434 -m 1 2 0 -m 1 3 0 -m 1 4 0 -m 2 1 0 -m 2 3 1.8181960943074 -m 2 4 0 -m 3 1 0 -m 3 2 2.02290154800773 -m 3 4 0 -m 4 1 0 -m 4 2 0 -m 4 3 0 -m 5 1 0 -m 1 5 0 -m 5 2 0 -m 2 5 0 -m 5 3 0 -m 3 5 0 -m 4 5 0 -m 5 4 0 -en 0.001 1 1.83290809268 -en 0.001 2 1.161030985567 -en 0.001 3 3.66914400056 -en 0.001 4 1.23640124358 -en 0.001 5 0.9132505 -em 0.020875 1 2 0 -em 0.020875 1 3 0 -em 0.020875 1 4 0 -em 0.020875 2 1 0 -em 0.020875 2 3 1.8181960943074 -em 0.020875 2 4 1.12888460726286 -em 0.020875 3 1 0 -em 0.020875 3 2 2.02290154800773 -em 0.020875 3 4 0.514005225416364 -em 0.020875 4 1 0 -em 0.020875 4 2 0.61034918826118 -em 0.020875 4 3 2.77081002950074 -em 0.042025 1 2 0 -em 0.042025 1 3 0.0447270935214584 -em 0.042025 1 4 0.00204350937063846 -em 0.042025 2 1 0 -em 0.042025 2 3 1.8181960943074 -em 0.042025 2 4 1.12888460726286 -em 0.042025 3 1 0.0340892941439601 -em 0.042025 3 2 2.02290154800773 -em 0.042025 3 4 0.514005225416364 -em 0.042025 4 1 0.00878072013784504 -em 0.042025 4 2 0.61034918826118 -em 0.042025 4 3 2.77081002950074 -en 0.104325 2 0.0402577179646081 -en 0.104325 3 0.192594746352967 -en 0.106325 3 8.73162876459514 -ej 0.106325 2 3 -em 0.106325 1 2 0 -em 0.106325 1 3 0.0177338314347154 -em 0.106325 1 4 0.00204350937063846 -em 0.106325 2 1 0 -em 0.106325 2 3 0 -em 0.106325 2 4 0 -em 0.106325 3 1 0.00723425109237692 -em 0.106325 3 2 0 -em 0.106325 3 4 0.193855714034029 -em 0.106325 4 1 0.00878072013784504 -em 0.106325 4 2 0 -em 0.106325 4 3 0.00771007640703268 -en 0.21195 5 0.1223036 -en 0.214175 5 0.194964 -en 0.267475 4 1.23640124358 -en 0.267475 5 0.194964 -ej 0.2675 5 4 -en 0.41955 1 0.158405393915496 -en 0.42155 1 0.299481445247702 -en 0.473075 4 0.0306317427630759 -en 0.475075 4 2.79429564470655 -en 0.480625 4 0.0872103733618782 -em 0.480625 1 2 0 -em 0.480625 1 3 0.0177338314347154 -em 0.480625 1 4 0.00204350937063846 -em 0.480625 2 1 0 -em 0.480625 2 3 0 -em 0.480625 2 4 0 -em 0.480625 3 1 0.00723425109237692 -em 0.480625 3 2 0 -em 0.480625 3 4 0.193855714034029 -em 0.480625 4 1 0.00878072013784504 -em 0.480625 4 2 0 -em 0.480625 4 3 0.00771007640703268 -en 0.482625 3 1.66920782430592 -ej 0.482625 4 3 -em 0.482625 1 2 0 -em 0.482625 1 3 0.241282075772286 -em 0.482625 1 4 0 -em 0.482625 2 1 0 -em 0.482625 2 3 0 -em 0.482625 2 4 0 -em 0.482625 3 1 0.0101771164248256 -em 0.482625 3 2 0 -em 0.482625 3 4 0 -em 0.482625 4 1 0 -em 0.482625 4 2 0 -em 0.482625 4 3 0 -en 1.5988 3 0.00336130452736601 -en 1.6008 3 1.47105091660349 -ej 1.6008 1 3 -em 1.6008 1 2 0 -em 1.6008 1 3 0 -em 1.6008 1 4 0 -em 1.6008 2 1 0 -em 1.6008 2 3 0 -em 1.6008 2 4 0 -em 1.6008 3 1 0 -em 1.6008 3 2 0 -em 1.6008 3 4 0 -em 1.6008 4 1 0 -em 1.6008 4 2 0 -em 1.6008 4 3 0

As a further assessment of the fit of the model, we plotted the observed and simulated site frequency spectrum (SFS), Figure 2-figure supplement 1. In general, the model fit is good, being poorest for singletons (too high) and high frequency derived sites (too low). This is likely due to effects of selection on the genome, which is not incorporated into the neutral demographic model. We note too, that this model was computed using only the allele counts from regions of the genome under weak/no selection as inferred from GERP scores, further explaining the reduced fit at these two site classes.

### Simulations of chimpanzee genetic data under neutrality and background selection

We used *msms* to perform coalescent simulations of chimpanzee demography. To simulate the effects of background selection (BGS) we modified the estimates of effective population size (*N_e_*) from the demographic model by multiplying them by a scaling factor, which represents the B score or effective reduction in *N_e_* due to BGS. 0.8, for example, reduces the *N_e_* and hence expected neutral diversity to 80% the level seen for neutral sites unlinked to regions under purifying selection (Key et al. 2016). We simulated non-genic regions with B=1, and genic regions with various strengths of BGS. We used B in the range 1-0.8, incremented by 0.025, with additional 0.0125 increments between 0.9 – 0.85. For neutral regions and for each B we simulated 25 million 2.0 kb loci. After processing and calculating allele frequencies, we performed *δ* and PBSnj genic enrichments as described previously. To estimate a BGS strength that best matched the observed *δ* genic enrichments, we performed a simple sum of squared differences, summed for each *δ* genic enrichment bin for each pairwise comparison.

### Estimating the number of extra eastern chimpanzee adaptive events

We use the structure of the block jack-knife to estimate the number of adaptive events that are needed to result in the PBSnj eastern tail genic enrichment being greater than that of central chimpanzees or generated by BGS. Recall that to estimate the error variance on the genic enrichment in each bin of PBSnj, we divided the genome into non-overlapping 200 kb blocks. For each block we have the number of genic and non-genic SNPs per bin of the PBSnj distribution.

For eastern chimps, there are 3475 genic SNPs contained within 842 blocks (i.e. 168 MB) in the PBSnj rhs tail i.e. with a PBSnj scaled length >= 0.8.

Of these, there are 468 blocks containing only 1 SNP i.e. 56% of blocks, 82 blocks with 10 or more outlier genic SNPs. i.e. 10% of blocks, with a maximum block count of 111 genic SNPs (Figure 4-figure supplement 1a).

We rank blocks by the number of genic SNPs that are outliers. Iterating over this sorted list we remove blocks and recalculate the enrichment for genic SNPs. We define matching as the number of iterations required to reduce the tail bin genic enrichment to below a target value. We chose to order by the number of eastern tail genic SNPs as this results in a monotonically decreasing genic enrichment with each block being removed.

### Haplotype/LD based tests of selection

*i*HS is the ratio of the extended haplotype homozygosities (EHH) of derived versus ancestral alleles at polymorphic loci. As EHH is measuring linkage disequilibrium (LD), a larger value indicates greater LD for the derived allele. Under neutrality, derived allele frequency is correlated with allele age, so *i*HS scores are standardised in bins of derived frequency. Standardised scores have a mean of 0 and standard deviation of 1. Outliers are typically defined as standardised *i*HS > 2. nSL is a related statistic, but it calculates haplotype homozygosity as the number of matching SNPs rather than genetic distance. This approach is less biased towards regions of low recombination and is reportedly more sensitive to the detection of soft sweeps (Ferrer-Admetlla et al. 2014). XP-EHH compares the homozygosity of focal haplotypes between populations and we only performed XP-EHH calculations for the sister taxa: central/eastern and Nigeria-Cameroon/western.

### Measures of conservation and effects on gene regulation

We used *phastCons* (Siepel et al. 2005) to infer highly conserved sites. We used the 20 mammalian multiz alignment to the human genome hg38, downloaded from UCSC (http://hgdownload.cse.ucsc.edu/goldenPath/hg38/multiz20way/maf/). To reduce the chance that polymorphism in chimpanzees affects inference of conservation, we removed both the chimp and bonobo reference genomes form these alignments. We estimated the phylogenetic models from fourfold degenerate (nonconserved model) and codon first position sites (conserved model). We then predicted base conservation scores and conserved fragments using the following options: --target-coverage 0.25 --expected-length 30. Resultant conserved elements covered 69.24% of the human exome, or an enrichment of 17.27. We note that although we attempted to remove the *Pan* branch from our estimates, it is impossible to completely avoid the use of these genomes, for example, when converting predicted conserved elements from hg38 to pantro2.1.4. These results have been deposited on Dryad (see Data availability).

We used regulomeDB (Boyle et al. 2012) to identify putatively regulatory role of genomic sites. Due to the close phylogenetical relationship between chimpanzees and humans, we argue that in lieu of any functional data for chimpanzees, inferred function from homologous positions in the human genome is a useful proxy for function in the chimpanzee genome. To obtain regulomeDB information for variable chimpanzee positions we used liftover to map SNP coordinates from pantro2.1.4 to hg19, keeping positions that reciprocally mapped to homologous chromosomes. Alan Boyle then kindly provided regulomeDB annotations for these positions. In regulomeDB, lower scores reflect higher confidence in regulatory function. We modified scores on the basis that scores 1a-f are given for positions that are human eQTLs, which we do not use as they refer to the specific allele change in humans rather than to the function of the site. Without eQTLS, scores 1a-c and 2a-c reflect the same biochemical signatures and location within transcription factor motifs. Thus, we combine these scores in to a new “high confident” regulatory function category. Our “non-regulatory” category includes positions with regulomeDB scores of 6 or 7, which have no evidence of being regulatory. We did not use sites with intermediate scores.

### Gene set enrichment analyses

We used GOWINDA (Kofler and Schlötterer 2012) to test for enrichments in Gene Ontology (GO) categories, which corrects for clustering and gene length biases. We used either GO categories or custom gene lists as candidate gene sets. GO categories for humans were obtained from the GO consortium (The Gene Ontology 2017, Ashburner et al. 2000), while gene sets were manually created from published sets of Viral Interaction Proteins (Enard et al. 2016) and a set of genes that are differentially expressed in CD4 cells after SIV infection in the natural SIV host vervet monkey but not in that non-natural host macaque (Svardal et al. 2017, Jacquelin et al. 2014, Jacquelin et al. 2009).

GOWINDA has an input file format which enables flexible usage of nonstandard gene sets. Genes are defined in a gtf file. We created a gtf from the ENSMBLE gene definitions, but restricted these to genes with clear 1-1 orthologs with humans. Our gtf file contained 16,198 of 17,530 protein coding genes. This gene set has been deposited on Dryad (see Data availability). Additional inputs are the PBSnj tail SNP set, and the background SNP set (of which the candidates are a subset). For all gene set enrichments, the background SNPs set was the full genome-wide set of genic variants for which PBSnj could be calculated.

GOWINDA was designed to reduce false positives that result from gene length bias (the probability of randomly containing an outlier SNP increases with gene length) and clustering of genes (such as paralogs) that share function. It achieves this by using resampling of background SNPs, which is the genome wide set of SNPs considered in a test. We use the -- mode *gene* switch. In this case, background SNPs are randomly sampled until the number of overlapping genes matches the total number of genes overlapping the PBSnj tail SNP set. Empirical *p-values* are estimated for each GO category, as the proportion of resamples which contain the same or greater number of genes than the PBSnj tail SNP set, per GO category (for each random background sample a pseudo *p-value* per GO category is also likewise calculated). FDR at each *p-value*, *p,* is then estimated as the number of observed *p-values* less than or equal to *p, R_obs_*, divided by the total number of resamples with a *p-value* less than *p R_exp_* i.e. FDR = *R_obs_* / *R_exp_*.

It is important to note that only genic background SNPs that are within the candidate set of genes (e.g. genes with GO definitions) are used in the random sampling. For the GO enrichment, after filtering for gene sets with at least 3 genes, the GO definition file contains definitions for 15649 genes, and 95% of genic background SNPs are used for resampling. This is important, as therefore GOWINDA cannot be used to directly test for enrichment in a single or small set of candidate gene sets. Providing one category, for example, would reduce the background SNP set to only those background SNPs in the genes in that category. Resampling can only ever return the same number of genes in this case. Thus, for VIPs and for the SIV gene set, we included an additional category, which is the full set of genes in the gtf file (“all gene set”). This has no effect on empirical *p-value* estimation. Its effect on FDR correction is limited as *R_obs_* is unchanged. For a candidate *p-value*, the all gene set will not be lower or equal to it unless the candidate *p-value* is itself 1. Thus *R_obs_* is unchanged. The effect on *R_exp_* is hard to determine, but for small empirical *p-values* should be proportionately small.

There are 98 VIP gene sets in (Enard et al. 2016), reduced to 53 when filtered for those containing at least 3 genes. For these and for the GO categories we used an FDR < 0.1 as a cut-off when discussing significant categories. There is only one SIV response genes set, so we only report the empirical *p-value* and treat *p-value* < 0.05 as significant. Note that this procedure does not allow the calculation of an FDR for the SIV set, nor over the family of tests (SIV gene set enrichment in all four subspecies) but we tested a strong *a* priori expectation that given the eastern PBSnj tail genes are enriched for viral immunity genes, this would be due to ververt SIV response genes. However, to estimate such an FDR, we used a resampling scheme: For each gene in the genome, we assign a weight, which is the proportion of SNPs in that gene compared to the genome as a whole. This is to correct for gene length bias. We make the intersect of all the SIV genes in each PBSnj tail. We then do weighted resampling from all genes in the genome to create sets of genes as large as the intersect set, and calculate an empirical *p-value* for each subspecies, as defined above. These empirical *p-values* are highly similar to those provided by GOWINDA, suggesting that our weighting scheme effectively controls for gene length bias. We then calculate the FDR for each empirical *p-value*, with *R_exp_* summed over all four subspecies.

### Natural Host SIV responsive genes underpin the eastern PBSnj tail genic enrichment

We wanted to test if selection on natural host SIV responsive genes could be the reason that Eastern chimpanzees exhibit the strongest signal of genetic adaptation. Our simple test is to hypothetically propose that if selection had not acted on the natural host SIV responsive genes then those genes would not contribute a SNP to the PBSnj eastern genic tail. Thus, we removed the genic tail SNPs from the 118 genes that are natural host SIV responsive and have SNPs in the outlier bin of the eastern PBS scores. However, we don’t remove the genic SNPs within these genes that are in any of the other subspecies. This means we will affect the eastern genic enrichment, but not the enrichment of other subspecies. We argue that this answers the question “what would the eastern genic enrichment be if selection had not acted on these genes in eastern chimpanzees”

### Statistics

To test enrichment in LD statistics, *phastCons* scores and regulomeDB scores we use random resampling tests. For a candidate set of SNPs sized *n*, we randomly draw the same number of genic SNPs. For LD statistics we calculated the mean score. For *phastCons* and regulomeDB we calculate the proportion of SNPs in a category. For the LD tests, all SNP scores are normalised so that genic SNPs have mean = 0 and sd = 1, within each bin of derived allele frequency. Thus, tail SNPs with a high score have a higher score compared to other genic SNPs with the same derived allele frequency.

For all resampling tests, *p-values* are estimated as 1 + *n* resamples >= observed (or <= observed as appropriate) / 1 + *n* resamples. Adding 1 to both the numerator and denominator ensures that resampling p-values do not equal 0, which is a downward biased estimate given finite resampling.

### Data availability

Data generated in the course of this investigation and relevant for the interpretation of the results presented here have been deposited with dryad: https://emea01.safelinks.protection.outlook.com/?url=https%3A%2F%2Fdatadryad.org%2Freview%3Fdoi%3Ddoi%3A10.5061%2Fdryad.2b3p518&data=02%7C01%7Cj.schmidt%40ucl.ac.uk%7C5b349af8089f4134a17b08d6a65d8526%7C1faf88fea9984c5b93c9210a11d9a5c2%7C0%7C0%7C636879317941617285&sdata=90ktL2z4I6XxuAX7F5qxMbWxdhoWEFpP8F3KoR8vJg0%3D&reserved=0

## Appendix 1

### Signed differences in derived allele frequency (*δ*) amongst human populations

We were interested in comparing the recent adaptive history of chimpanzees and humans. Previously (Coop et al. 2009) found that that those SNPs with the greatest allele frequency difference between populations of modern humans were enriched for genic variants. Subsequent work presented by (Hernandez et al. 2011) and (Key et al. 2016) replicated these findings. To present consistent analyses and make more specific comparisons with chimpanzees, we replicated the analyses of signed differences in derived allele frequency (*δ*) in three human populations: Yoruba in Ibadan, Nigeria (YRI); Japanese in Tokyo, Japan (JPT); British in England and Scotland (GBR). We choose JPT and GBR because their pairwise F_ST_ is 0.10 i.e. approximately the same as for eastern and central chimpanzees. The YRI- vs. non-African pairwise comparisons have amongst the largest F_ST_ values of all comparisons among 1000 Genomes populations, ~0.15 (data not shown). By down sampling each population to n =10 or n = 20 individuals, we can also assess the impact of sample size, considering that the range of chimpanzee samples is 10 −19. We used genotype data from the 1000 genomes phase III (Genomes Project et al. 2015), without any filtering of genotypes. We used the annotated ancestral allele in this same dataset (which are derived themselves from EPO alignments) to polarise derived allele frequencies.

We find that sample size has moderate effects on the determination of *δ* tail bin genic enrichment, except for GBR vs. JPT, which have few SNPs with a large frequency difference, and for which the high resolution afforded with sample n=91 is required to ascertain a significant tail bin enrichment (Figure S1a).

When comparing the 95% confidence intervals of tail bin genic enrichments we find, consistent with previous results, that *δ* tail bin genic enrichments are symmetrical for human populations when ancient DNA information is not incorporated (Key et al. 2016). This finding is consistent across all sample sizes (Figure S1a), suggesting that sample size is not a contributor to the stronger genic enrichment in central vs. eastern chimpanzees.

To further explore asymmetry among the genic enrichment in the two tails of *δ*, we repeat the calculation of tail bin genic enrichment log_2_ ratios as in Main Text Figure 2. In nearly all cases the enrichment is symmetric (Figure S1b). The only significant asymmetry is for an increased genic enrichment in Yoruba vs. Japanese when the sample size is 91. We note that despite being significant, this asymmetry is only half that observed in the comparison between eastern chimpanzees and central chimpanzees. No human comparison thus shows signatures that compare to those between eastern and central chimpanzees.

## Appendix 2

### Estimating the strength of background selection required to explain *δ* bin genic enrichments

Previously it has been shown that background selection (BGS) can result in genic enrichment in sites with large frequency differences between populations (Coop et al. 2009, Hernandez et al. 2011, Key et al. 2016). To find the strength of BGS (measured as a B score, a fraction of the expected neutral diversity) that could explain genic enrichments observed in chimpanzees, we simulated 25 million 2kb loci for non-genic (B = 1) and genic regions. For genic regions we used a range of B (1 – 0.8) in 0.025 steps, except between 0.9 – 0.85 for which we used a step size of 0.0125. While the strength of purifying and background selection varies among genes and genomic regions, this global (average) inference allows us to make comparisons at the genome scale. We used the sum of squared differences between simulated and observed genic enrichments for *δ* bins to ascertain which B provides the best fit to observed genic enrichments.

We find that when fitting all *δ* bins for all pairwise *δ,* the best fit is provided by B = 0.888 (Supplementary file 4), which indicates a reduction in neutral diversity levels of 11% in genes when compared with non-genes.

We repeated this exercise using data on the 12 *δ* tail bins alone. Doing so allows us to infer the strength of BGS required to fully explain the genic enrichment in the most highly differentiated SNPs, which likely harbour targets of positive selection. While assuming no influence of positive selection is unrealistic, this allows us to explore whether background selection alone could, in theory, explain our observations. The best fitting B in this case is 0.863, or a 14% reduction in genic diversity levels due to BGS (Supplementary file 4).

In contrast, by excluding the 12 *δ* tail bins, the fit of observed to simulated genic enrichments is less likely to be reduced due to the influence of targets of positive selection. The best fitting B in this case is 0.925 (Supplementary file 4).

Comparing the relative order of magnitudes of the Sum of Squares shows that the worst fit of simulated and observed genic enrichment are seen when attempting to fit all *δ* bins. This is an indication that BGS is not the only force affecting drift and diversity levels in genes, and combined with the observation that the best fit is B=0.925 when *δ* tail bins are excluded suggests positive selection is contributing to the genic enrichments in the *δ* tails.

We also checked if a greater genic enrichment in eastern vs. central chimpanzees is expected given the demographic history of chimpanzees and/or the effects of BGS. For each value of B we modelled above, we also calculated the log_2_ ratio of the eastern and central *δ* tail bin.

No value of B in the range 1 – 0.8 results in an asymmetry in genic enrichment between eastern and central chimpanzees as great as that observed in the genomic data (max B = 0.850, 0.103; observed 0.34). No large asymmetry is generated under the demographic model without BGS (B =1). Both results suggest that no combination of BGS strength can produce the difference in eastern and central *δ* tail bin genic enrichment observed.

## Appendix 3

### Evidence for, and explanatory power of, differing strengths of BGS amongst chimpanzees

We note first the evidence suggesting that background selection varies little among the great apes despite their large differences in *N_e_* (Nam et al. 2017) and despite the stronger purifying selection in larger *N_e_* subspecies (Bataillon et al. 2015). Background selection is expected to reduce diversity in genic regions more than in non-genic ones by removing variants linked to deleterious alleles, but the action of this type of selection appears independent of *N_e_* (Nam et al. 2017). Background selection is instead determined by the distribution of fitness effects for deleterious alleles, which is likely similar among the great apes owing to their generally conserved gene location and function (Nam et al. 2017). Further, simulations show that the rate of selective sweeps explains the larger reduction of diversity around genes in species with larger *N_e_* (Nam et al. 2017). Thus, the diversity reducing effect of background selection should be the same across all four chimpanzee sub-species. We tested this comparing the levels of scaled neutral diversity (π / divergence to macaque) between chimpanzee sub-species as a function of the distance to the nearest gene (normalized to the lowest diversity seen for each of the sub-species, Appendix 3-figure 1a). We confirmed that the relative reduction in neutral variation linked to genes is the same across sub-species (both Appendix 3-figure 1a), and that the nucleotide distance from genes at which neutral diversity reaches equilibrium is also similar. In addition, we find that the average genomic diversity in central, eastern and Nigeria-Cameroon chimpanzees has similar dependency on recombination rate and density of functional features (gene coding and gene untranslated sequences and non-coding conserved elements (Appendix 3-figure 1b) suggesting yet again that background selection is comparable among them. Note however that functional categories appear to be worse predictors of diversity levels in western chimpanzees than the other subspecies (95% CI of the bootstrap distributions of *rho*, the partial *spearmans* correlation controlling from recombination rate, do not overlap). We have not investigated this further, but it is possibly due to the fact that the genetic map is based on a sample of western chimpanzees, is therefore most accurate for this subspecies with the effect of smaller residuals in the regression of diversity on recombination rate.

Lastly, we turn to a population genetic statistical model able to estimate the reduction in neutral diversity due to background selection(Corbett-Detig, Hartl, and Sackton 2015). Full details for this model are given in Corbett-Detig *et. al.* (2015), but we briefly recapitulate the main points. The effect of BGS is estimated as the population scaled mutation rate (4*N_e_μ*, θ) scaled by a parameter G, that models BGS as a local reduction in *N_e_* with G allowed to vary in windows along the genome in proportion to the per window fraction of functional sites. The effect of selective sweeps or Hitch Hiking (HH) is also estimated as θ divided by the population scaled rate of sweeps, 2*Nv*. Following the implementation of this model by Corbett-Detig *et. al.* (2015) we calculated average neutral diversity in 500 kb windows and used the number of bp in exons for functional density. We ran the compute_gk package on this data to estimate the effects of linked selection, as described by Corbett-Detig *et. al.* (2015). Using this model, we estimated the same reduction in neutral diversity in each chimpanzee subspecies (11% reduction in the highest likelihood model; Supplementary file 6), indicating equivalent levels of background selection among sub-species.

Despite there being no evidence to suggest that there are differences in the strength of BGS between eastern and central chimpanzees, it is useful to determine if such a putative asymmetry in BGS strength could lead to eastern chimpanzees having a greater tail bin genic enrichment than central chimpanzees. To investigate this possibility, we performed simulations of BGS, where the strength of BGS was stronger in eastern chimpanzees (B = 0.825) than in all other chimpanzees (B range: 0.900 – 0.850). We find that while a greater eastern B marginally increases the relative magnitude of eastern chimpanzee *δ* tail bin genic enrichment, none of the simulated ratios are within the 95% of the eastern vs. central *δ* tail bin log_2_ ratio. This further reinforces stronger BGS in eastern than in other chimpanzees would result in differences in *δ* tail bin genic enrichment.

## Appendix 4

### Population branch statistics

In the two-population case (Pop A, Pop B), a “scan” for targets of population selection can be performed by identifying outliers – e.g. the top 5% of sites – in the genome wide distribution of per site pairwise F_ST_ values, if one assumes that these outliers are likely enriched for true targets of positive selection (the empirical distribution could also be compared to simulated values). As pairwise F_ST_ is a summary of the joint site frequency spectrum (SFS) of two populations, these outliers are the sites with the greatest site (allele) frequency difference. If one considers this as an unrooted two population tree (i.e. a straight line), outliers are simply those sites with the longest branch lengths. The problem of course is that there is no directionality to F_ST_, but one assumes that the population with the highest derived allele frequency is the one in which selection has acted.

The Population Branch Statistic (PBS), introduced by (Yi et al. 2010) in their study that identified *EPAS1* as under selection in Tibetans, extends the pairwise F_ST_ case by the addition of a third population, Pop C. PBS is a function of the three possible pairwise F_ST_ values amongst three populations (AB, AC, and BC). As in the two-population case, there is only one unrooted tree relating three populations, with each population connected to the central node. Therefore, each population can be assigned a unique branch length or PBS value (Appendix 4-figure 1a). The branch length is indicative of the population specific change in allele frequency, and targets of positive selection can be identified as outliers. Thus, PBS overcomes the issue of assigning directionality to allele frequency differences between populations, although with the assumption that selection occurs in one branch only.

We wanted to analyse the joint frequency spectrum of the four chimpanzee subspecies, and used PBS as an inspiration to develop a new statistic, PBSnj. We analyse a simple four population model, with two groups of sister taxa A,B and C,D sharing a common ancestor AB,CD. Split times for AB,CD, A,B and C,D are 0.2, 0.1, 0.1 scaled time units respectively, and population size is 10e^3^ throughout. We performed 2 million simulations of a 2kb locus, with mutation rate = 1.2e^-8^ and recombination rate = 0.96e^-8^, and sampling 50 chromosomes per population. The msms (Ewing and Hermisson 2010) command line used is:

msms 200 1 -t 0.96048 -r 0.768384 -I 4 50 50 50 50 -n 1 1 -n 2 1 -n 3 1 -n 4 1 -en 0.1 2 1 -en 0.1 4 1 -ej 0.1 1 2 -ej 0.1 3 4 -en 0.2 4 1 -ej 0.2 2 4.

We take A as the focal population. There are three possible combinations of F_ST_ values to calculate the branch length leading to population A: ABC, ABD, and ACD, denoted PBS_ABC_ *etc*. We note that in the Tibetan PBS example, populations were chosen so that one was clearly ancestral: Danish is the outgroup to Tibetan and Han. This highlights that while the underlying tree is unrooted and the Tibetan branch represents allele frequency change since their split with Han Chinese, in reality the Danish branch is a compound branch length combining branches leading from the basal Eurasian common ancestor to the Danish and the basal Eurasian common ancestor to the common ancestor of Tibetans and Han Chinese. In this sense, the Danish PBS branch would not represent population specific selection events *per se*, and its length is not an indication of selection events in the Danish. This indicates that the ability of PBS to truly distinguish population specific allele frequency changes is dependent on the configuration of populations included in its computation. To show this is true, we plot the rank correlations of the three different PBS statistics possible for PopA. PBS_ABC_ and PBS_ABD_ are highly correlated (*spearman’s rho* = 0.82, Appendix 4-figure 1b) but both are poorly correlated with PBS_ACD_, which is a compound branch length in our model (*spearman’s rho* = 0.46 and 0.47; PBS_ACD_ vs. PBS_ABC_ and PBS_ABD_, Appendix 4-figure 1b).

That PBS_ABC_ and PBS_ABD_ are not perfectly correlated indicates that each contains independent information in delimiting the branch length of PopA, and illustrates the motivation in producing a statistic that draws upon the full four population F_ST_ matrix.

In deriving this statistic, we note that PBS is just a simple algebraic function of the matrix of pairwise F_ST_ values. To find PBS_ABC_, for example: PBS_ABC_ = (distanceAB + distanceAC – distanceBC)/2. An alternative method for finding distances in a phylogeny is the Neighbor-Joining algorithm (NJ) (Saitou and Nei 1987). Without giving the full details, NJ proceeds by calculating a *Q* matrix from the input distance matrix, creating a node by grouping the two taxa with the smallest *Q*, and re-calculating distances with respect to the new node. In this sense, branch lengths are a by-product of the NJ procedure, but nonetheless, by recording these branch lengths for each SNP NJ tree across the genome, we can generate a distribution of branch lengths analogous to PBS. For this reason, we name this proposed statistic PBSnj. While the details and actual distances calculated differ, PBS and PBSnj both define a distance for each branch in a tree, and the correlation between three-population PBS and PBSnj branch lengths suggests that these two methods are near identical in their results (spearman’s *rho* = 0.995). Extending PBS to more than three populations require fixing a topology. In the four population case, branch length A could be calculated as: PBS_ABCD_ = (PBS_ABC_ + PBS_ABD_) / 2, but this assumes that the tree at each site follows the species tree ((A,B), (C,D)). It also “hides” the presence of an internal branch implicit in a bifurcating four taxa tree. While more complicated sets of algebraic functions could be combined to solve this or other conundrums, it is enough to point out that nj does not assume a topology (it is after all a topology finder) and that its algebraic rules are consistent no matter the number of taxa, the only change being the number of repetitions of the algorithm. Thus, we conclude that PBSnj is the more natural method to use. Lastly, while we have not considered it here, in theory PBSnj is extendable to any number of taxa.

We also do not consider the internal branch as in this investigation we are only interested in the selection pressures that differentiate extant populations of chimpanzees. Furthermore, interpretation of the direction of the internal branch in the four taxa case relies again on assuming that the derived allele is the target of selection.

The schematic for calculating PBSnj is as follows:

1. for each site, calculate the full F_ST_ matrix.
2. apply the Neighbor-Joining algorithm on the F_ST_ matrix, i.e. generate a nj-tree.
3. for each site, record the branch length for each taxa in the nj-tree.

Following the original description of PBS (Yi et al. 2010), we transform the F_ST_ values into units of drift time: -ln(1-F_ST_). As this is undefined for F_ST_ == 1, we substitute F_ST_ == 1 for the next lowest possible pairwise F_ST_ value. So that branch lengths exhibit the same range, following (Malaspinas et al. 2016) we standardised branch lengths by the total length of the tree, e.g. PBSnj_A_scaled_ = PBSnj_A_ / (1 + PBSnj_A_+ PBSnj_B_ + PBSnj_C_ + PBSnj_D_ + PBSnj_INTERNAL_). Lastly to perform genic enrichment tests analogous to derived allele frequency difference we re-scale so that values are within the range 0-1. This implies values of PBSnj >= 0.8 (which we use as our cut-off or PBSnj genic tail bin) are those equal to 80% or more of the max possible values of PBSnj, are not a quantile cut-off and can therefore contain a differing number of sites per taxa.

As a simple illustration of the effectiveness of PBSnj to identify population specific changes in allele frequency, we asked how well the statistics identify Pop A specific allele frequency change. We plot the derived allele frequency in each of Pops A-D, for those sites for which PBSnj_A_scaled_ >= 0.8 (Appendix 4-figure 1c). As a comparison, we do the same for PBS_ABC_, PBS_ABD_ and PBS_ACD_. PBSnj_A_ clearly delineates those sites specifically differentiated in Pop A. PBS_ACD_ is the worst test statistic, as Pop B allele frequencies are nearly uniformly distributed in the range of 0-1 despite these sites being identified as pop A outliers. PBS_ABC_ and PBS_ABD_ offer a substantial improvement, but note there is a tendency for a more uniform distribution of allele frequencies in the population not included in the calculation of PBS_ABC_ and PBS_ABD_ (D and C respectively). Note too, the point masses near 0 for Pop A, and near 1 for Pops B-D in PBSnj_A_scaled_ which represent those sites where PopA has a very low derived allele frequency i.e. are ancestral allele outliers.

## Appendix 5

### The relationship between divergence times and N_e_ and the effects of BGS

Demography varies greatly amongst the chimpanzee sub-species, with a wide range of pairwise divergence times and effective population sizes *N_e_* (see Main Figure 1). This means that the total genetic drift between, for example, Western and Central chimpanzees is much greater than that between Central and Eastern chimpanzees. It is unknown how these differences in drift times effects genic enrichments in bins of either the signed differences in derived allele frequency (*δ*) or PBSnj. To explore this, we again used a simple four population model (described Appendix 4). To model the effect of background selection (BGS) we can scale *N_e_* by a value B, such that *B*= 0.9, for example, represents BGS that reduces linked neutral diversity by 10%. Genic regions were simulated using B = 0.9. We also allowed either the basal split time, or the split time of pop1 and pop2 to increase, therefore widening the range of divergence times.

For each scenario, we simulated 50 chromosomes per deme for a 2kb locus, for 2 million replicates, using a mutation rate of 1.2 × 10^-8^ and recombination rate of 0.96 × 10^-8^.

### The effect of divergence time and N_e_ on δ tail bin SNP n

Increasing the divergence time increases the number of SNPs in both genic and non-genic tail bins, as is expected due to a greater variance in allele frequency due to genetic drift. While intuitive, it is important to demonstrate as it shows that the number of SNPs in tail bin is not itself an indication of the statistical support for selection. Increasing divergence time also reduces the genic:non-genic SNP n ratio: from ~1.5 at time = 0.1 down to ~1.08 at time = 0.4.

Changes in *N_e_* also greatly affect the number of *δ* tail bin SNPs. We varied the simulated PopB:PopA ratio to be either 0.9, 0.5 or 0.1. On this time scale, an *N_e_* ratio of 0.9 has a modest impact on the number of Pop2 tail bin SNPs. However, ratios of 0.5 and 0.1 result in a dramatic increase in both Pop2 genic and non-genic tail bin SNPs. Of course, this mirrors the result of increasing divergence time – for the same evolutionary time, lower *N_e_* results in greater drift. What is also apparent is that that the lowered Pop2 *N_e_* also results in an increase in the Pop1 *δ* tail bin counts. When Pop2 *N_e_* = 0.1, the ratio of genic to non-genic *δ* tail SNPs is ~ 1 for both populations. We posit that these two factors – increased divergence and lower effective population size – explain the lower genic enrichments seen for *δ* calculated with western chimpanzees. A secondary point is the implication that the genic enrichment produced by a given strength of BGS decreases with drift time.

### The effect of divergence time and N_e_ on PBSnj tail bin genic enrichment

Divergence time and *N_e_* impact *δ* tail genic enrichment of both populations. This is because it conflates allele frequency change occurring in two populations. In contrast, PBSnj is able to determine the allele frequency change that occurs specifically in one branch of a phylogeny. To show the effect of *N_e_* on the PBSnj tail genic enrichment, we plot the genic enrichment assuming BGS of B =0.9 but varying the *N_e_* of Pop2 (Appendix 5-figure 1). Given a relative *N_e_* = 1, the genic enrichment in the PBSnj tail bin = 1.20, and there is no effect in reducing the *N_e_* to 0.9 (Appendix 5-figure 1). Below *N_e_* = 0.9, the genic enrichment drops precipitously to 1.16 for *N_e_* = 0.5 and 1.06 for *N_e_* = 0.1. We suggest that this shows that BGS has a greater impact when divergence times are shorter and *N_e_* relatively large, that is when most of the variation between lineages is still segregating. Longer divergence times and lower Ne results in a greater number of fixed differences between lineages, and BGS does not impact the divergence in genic regions to the extent that it reduces diversity and distorts the SFS of segregating variation.

This result is the motivation for comparing only central and eastern chimpanzee PBSnj tail genic enrichments. Not only is their pairwise divergence time the lowest amongst the chimpanzees, but given their relative *N_e_*, we would not expect *N_e_* to be the reason that eastern chimpanzees exhibit a greater PBSnj tail genic enrichment. Indeed, simulations recapitulating the demographic history of chimpanzees suggest that BGS produces equal genic enrichments for eastern and central chimpanzees. As well as expecting a similar level of drift in each of their branches, given a constant rate of adaptive evolution, we would also expect a similar number of adaptive events to contribute to the genic enrichment.

## Appendix 6

### Estimating the strength of background selection required to explain PBSnj tail genic enrichments in chimpanzees

We also determined how background selection can affect the PBSnj statistic amongst chimpanzees and found that they each have a unique value of B which best explains their PBSnj tail bin genic enrichment. We explain this by positive selection differentially influencing the tail of each species, as we do not expect (Nam et al. 2017) or observe differences in the effects of background selection across species. However, only eastern chimpanzees require a B stronger than 0.888 to achieve the observed PBSnj tail genic enrichment.

Of critical importance for interpreting the greater PBSnj tail genic enrichment for eastern compared to that for central chimpanzees is the observation that across all values of B tested, simulated genic enrichments are approximately identical for these two subspecies (Figure 4-supplement 2b). Thus, demography and BGS should not produce the observed pattern. Results from our generalised four population model also indicate that the relatively small difference between eastern and central *N_e_* are also not a likely explanation. In fact, such differences should result in a higher enrichment for central chimpanzees who have the larger *N_e_*. We again posit that this is evidence for a greater rate of adaptive events along the eastern branch than that for the central branch.

In contrast, for any given strength of BGS, the simulated eastern and central genic enrichments are always greater than those of western and Nigeria-Cameroon chimpanzees. Our explanation for this is as follows: most tail SNPs for Nigeria-Cameroon, and especially for western chimpanzees, are actually fixed differences to all other chimpanzees. On the other hand, most eastern and central PBSnj tail SNPs are polymorphisms shared between these two sub-species. In addition, results from our general four population model, indicate that by increasing the lineage specific drift by increasing divergence time and/or decreasing *N_e_*, the genic enrichment caused by BGS decreases. Again, we suggest that this indicates that BGS is more important for polymorphism than divergence. Finally, we conclude that only the eastern vs. central PBSnj tail bin comparison is informative in judging the significance of the eastern PBSnj tail genic enrichment or the likelihood that this can be explained by BGS.

## Appendix 7

### Demography and the evidence of positive selection in central chimpanzees

We use this section to also discuss the factors that could blunt the evidence for positive selection in central chimpanzees. Population structure within the sampled central chimpanzees could reduce the apparent number of highly differentiated alleles. Fixed beneficial alleles in two divergent central chimpanzee populations would, for example, look to be segregating at intermediate frequencies if these were both sampled equally. Population structure within chimpanzee subspecies has been extensively analysed by both (Prado-Martinez et al. 2013) and (de Manuel et al. 2016). de Manuel et. al. (2016) present the results of numerous analyses within their supplementary material, including results from sNMF (Frichot et al. 2014), fineSTRUCTURE (Lawson et al. 2012), and ADMIXTURE (Alexander, Novembre, and Lange 2009) (respectively figures S11, S13 and S14 in de Manuel et. al. (2016)). For each of these analyses, there is less structuring of the sampled central chimpanzees compared to the sampled eastern chimpanzees, which in contrast often appear as a cline of variation from Tanzania and the south of the Democratic Republic of the Congo (DRC) through to Uganda and northern DRC. This suggests that unaccounted-for population structure is not a reason for weaker genic enrichment of differentiated alleles in central chimpanzees.

Another possible blunting mechanism is gene flow. When simulating neutral evolution and BGS, we used coalescent simulations using demographic parameters previously described in de Manuel et. al. (2016). This model includes inferred gene flow amongst Pan lineages, including that of the bonobo – central chimpanzee introgression. However, what these simulations do not address is the possibility that alleles selected in central chimpanzees were constantly stopped from reaching fixation due to the introduction of bonobo alleles until cessation of this gene flow ~40 kya. We note first, that gene flow is a general barrier to local adaptation, and that the rate of migration and strength of selection are the two key parameters determining the likelihood of reaching fixation. If gene flow into central chimpanzees was too great or selection too weak, then this could reduce the genic enrichment in population specific, highly differentiated alleles – but this would reflect the biological reality of reduced local adaptation in this subspecies. Secondly, we highlight that while bonobo introgression into central chimpanzees did occur, the scale of this gene flow is dwarfed by the ongoing, and near symmetrical, gene flow between central and eastern chimpanzees (migration into central chimpanzees from bonobo was ~ 1.6% of the ongoing rate from eastern chimpanzees, and ~ 1.9% of the ongoing rate of migration from central into eastern chimpanzees). These rates of migration would pose a greater barrier to adaptive population differentiation, and the signal in eastern chimpanzees is identified despite this.

## Acknowledgements

Acknowledgments. We thank: Fabrizio Mafessoni, Linda Vigilant, Mimi Arandjelovic, Paolo Gratton, Hjalmar Kühl and Lauren White of the Max Planck Institute for Evolutionary Anthropology for helpful discussions and/or comments on the manuscript; Alan Boyle for providing regulomeDB scores; Hannes Svardal for extensive discussions and comments on the manuscript. This work is (partly) funded by the NIHR GOSH BRC. The views expressed are those of the author(s) and not necessarily those of the NHS, the NIHR or the Department of Health.

## Fihure legends

**Figure 2-figure supplement 1:**
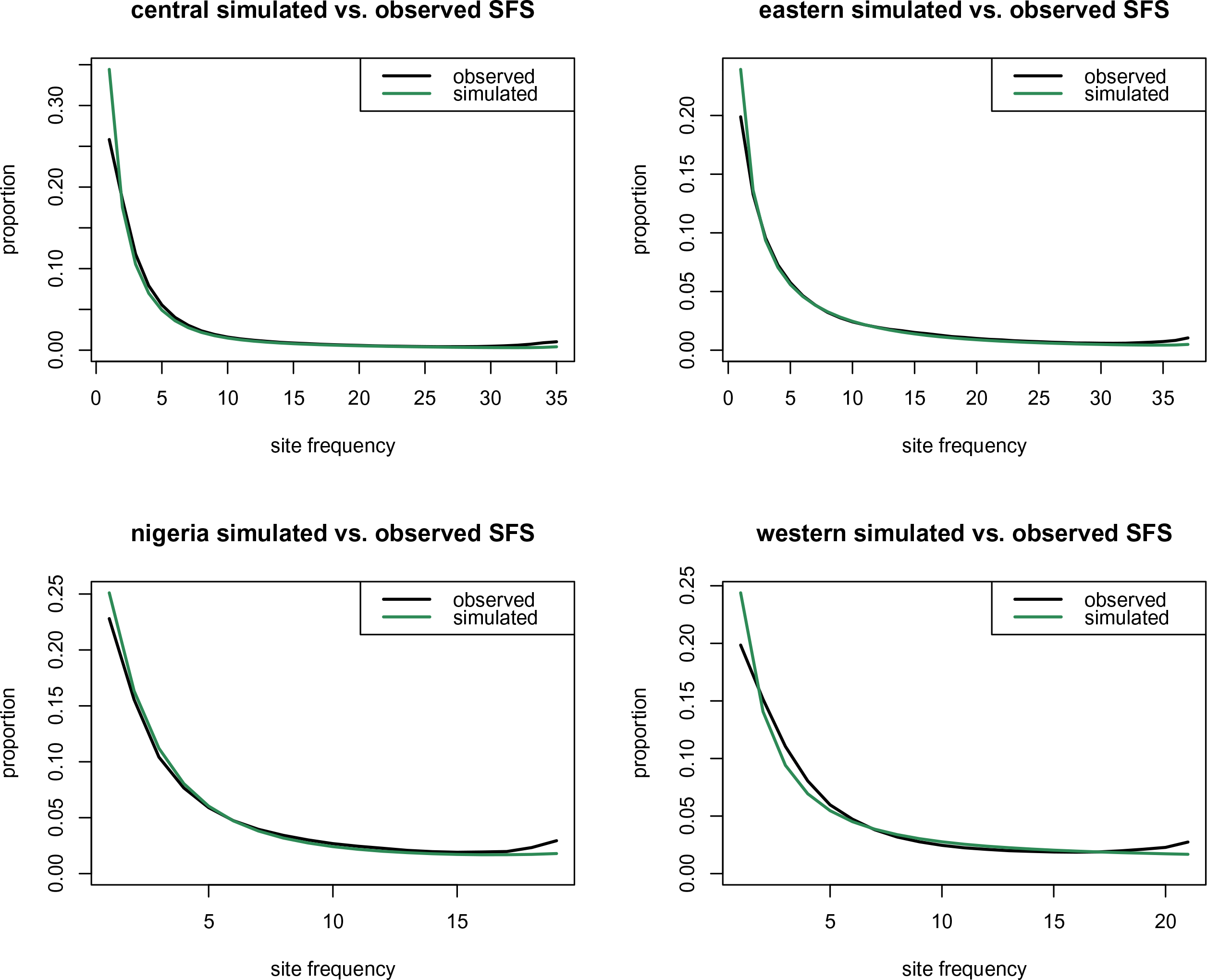
Observed and Simulated Site Frequency Spectra. We plot the Site Frequency Spectrum (SFS) for each chimpanzee subspecies. X axes: derived allele count. Y axes: proportion. Black: observed. Green: simulated. Simulated counts come from 25 million 2kb loci simulated with *msms*, using the chimpanzee demography specified in Methods.

**Figure 3-figure supplement 1:**
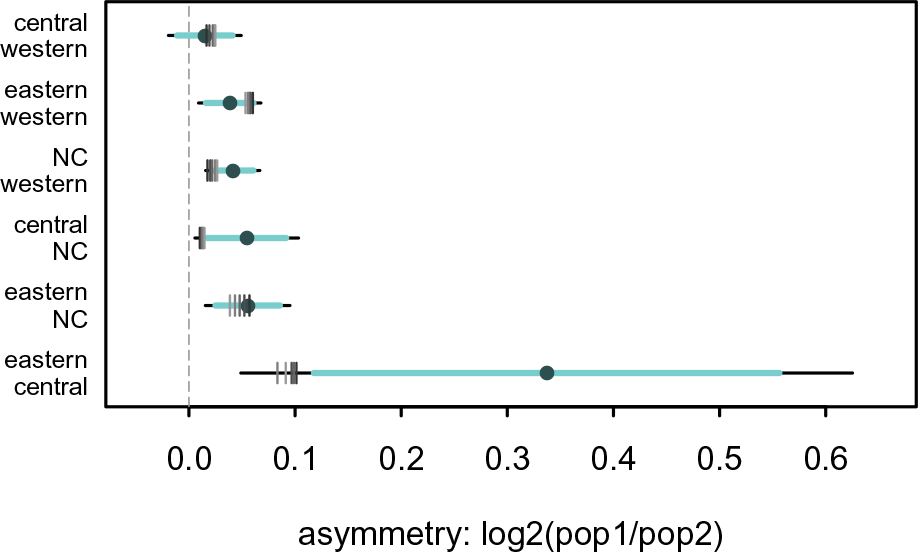
Stronger eastern BGS does not result in observed levels of δ tail bin genic enrichment asymmetry. The asymmetry of the genic enrichments in the *δ* tails is measured by taking their log_2_ ratio, thus 0 indicates a symmetric enrichment (equal enrichment in both *δ* tails). We created coalescent simulations in which the strength of BGS was greater in eastern chimpanzees than other subspecies. For eastern chimpanzees we chose a fixed B = 0.825, as this B provided the best fit the eastern *δ* tail genic enrichment. All other subspecies had the same B, in the range of 0.900 – 0.850. A larger difference in B between subspecies results in a slight increase in asymmetry, but none of the simulated differences in BGS result in the observed asymmetry. Point = observed asymmetry. Horizontal lines represent confidence intervals estimated by 200kb weighted block jackknife (light = 95%, black = 99%, i.e. alpha = 0.05 or 0.01 for a two-tailed test). Grey vertical marks represent the *δ* tail asymmetry in simulations, under increasing levels of difference in background selection between eastern and other chimpanzees: lightest to darkest shades: All B_eastern_ = 0.825; B_others_ = 0.850, 0.863, 0.850, 0.888, 0.900.

**Figure 4-figure supplement 1:**
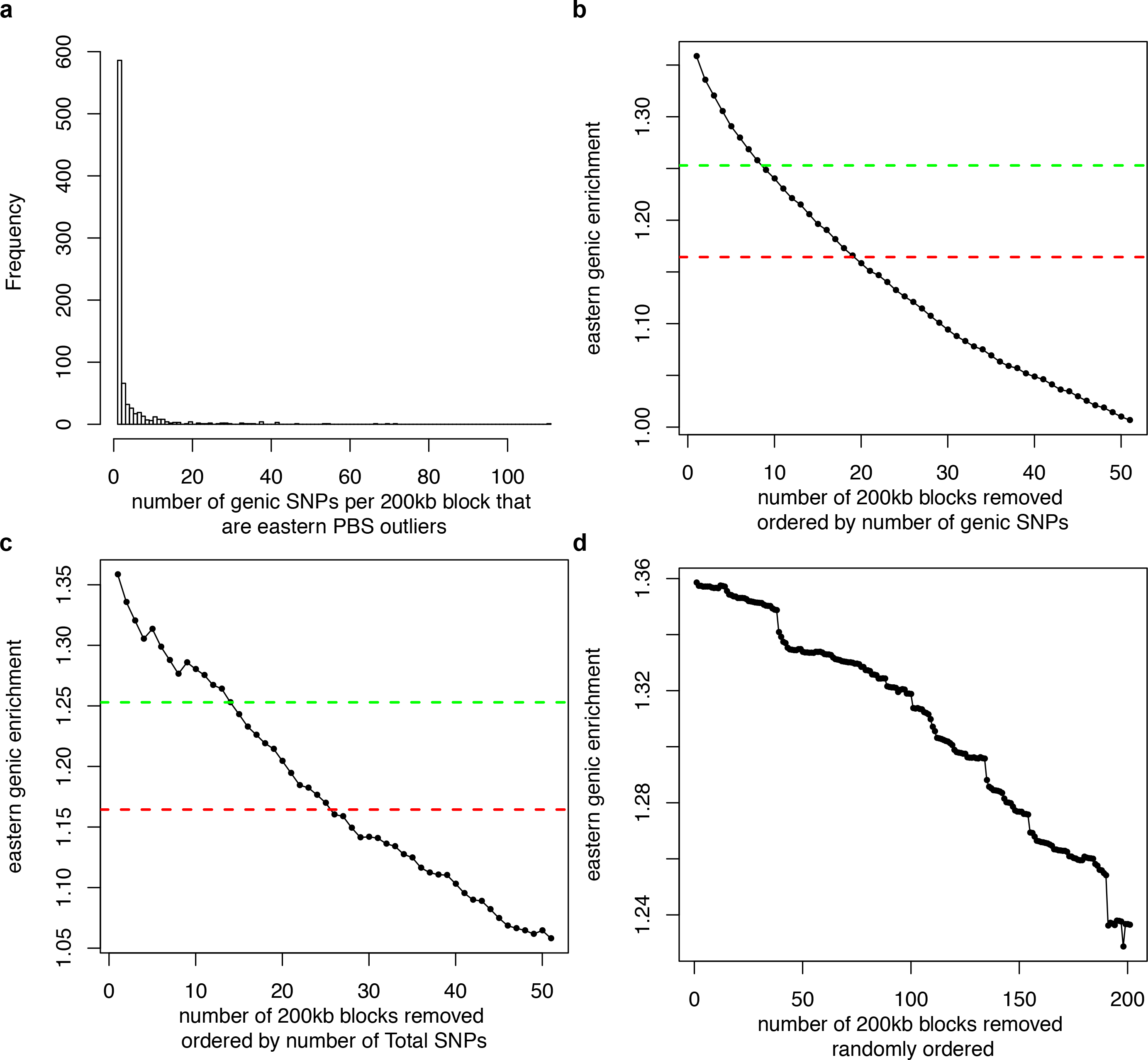
Number of adaptive events in eastern chimpanzees. **a,** Most 200kb blocks contain few PBSnj eastern outlier SNPs, but there is an extended right hand tail. **b,** we ranked blocks by the number of PBSnj eastern tail SNPs, then iteratively removed outlier genic SNPs. This results in a monotonically decreasing genic enrichment, and the removal of eight blocks is required to reduce the genic enrichment of the PBSnj eastern tail to overlap the 95% CI of the PBSnj central tail, and 19 blocks to reduce it below the level of the point estimate of the central PBSnj tail. We could alternatively order the windows by the total number of outlier SNPs i.e. without regard to genic vs. non-genic. Doing so increases our estimated range of sweeps to 15-26. But we note that the genic enrichment does monotonically decrease with block removal (**c**). This is partly due to the arbitrary nature of the definition of genic, as it implies that there are some 200 kb blocks that have more non-genic than genic outlier SNPs contained within them, and this may very well change if the definition of genic was changed from transcription start and end sites +-2kb. (**d**) Lastly, we randomly shuffled the removal order of the 200-kb blocks. We did so for 1000 random shuffles of the block order (A single random shuffle is shown). We find that the median number of blocks (i.e. sweeps) across random shuffles is 165 to match the upper 95% CI of the central chimpanzee estimate (middle 90% quantile range 114-221; min = 78, max = 278) increased to 273 to match the central chimpanzee point estimate of genic enrichment (middle 90% quantile range 214-329; min = 162, max = 381). Such a procedure is likely an overestimate, as most of the removal steps are those removing 1 to 9 genic outlier SNPs (panel a), resulting in minimal reduction of the genic enrichment.

**Figure 4-figure supplement 2:**
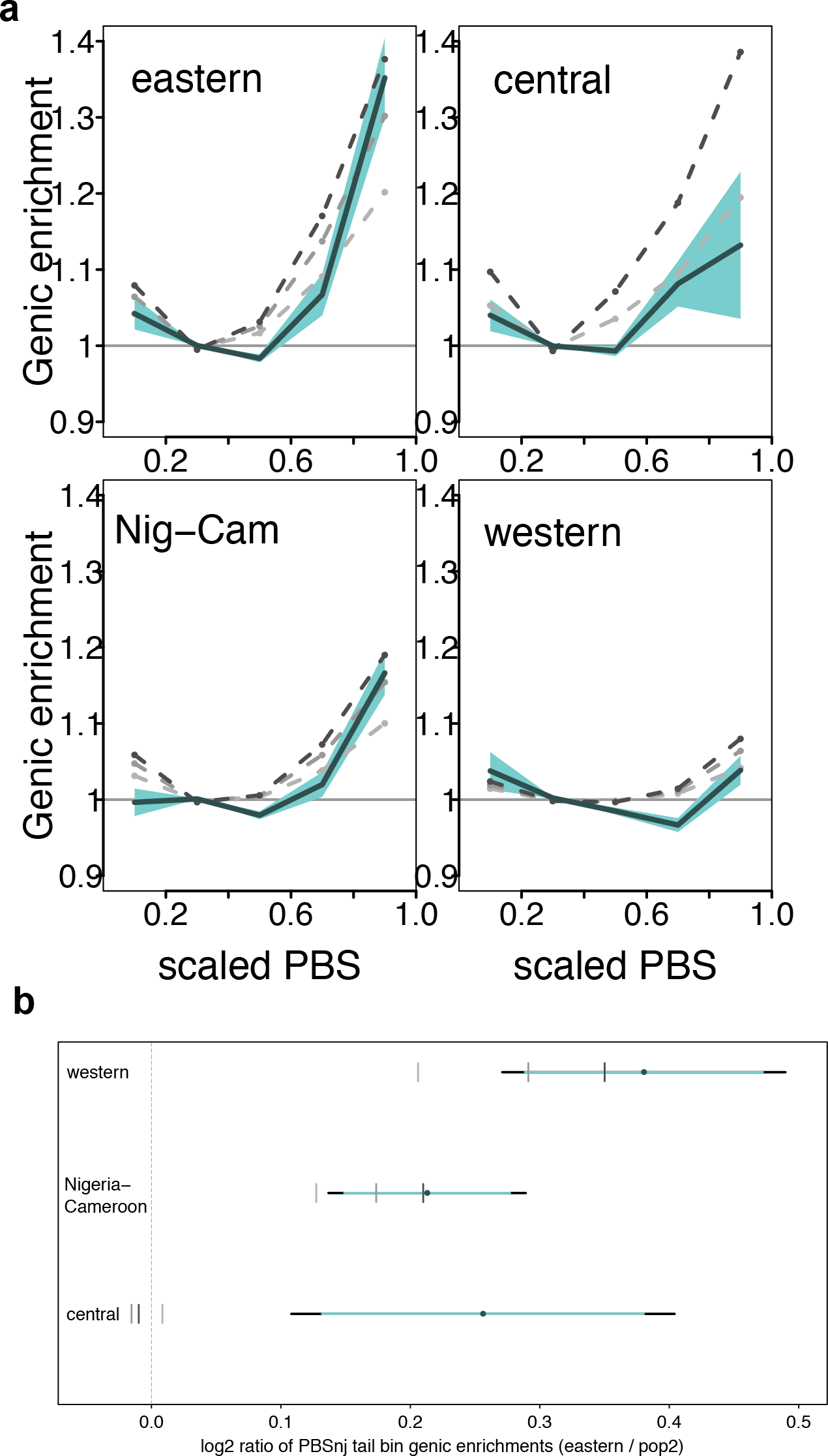
Scaled PBSnj bin genic enrichment for all chimpanzee subspecies. **a,** X-axes: PBS scaled to take values in the range 0 −1, per subspecies. Y-axes: Genic enrichment computed as described in Fig. 2. Shading represents the 95% CI (i.e. alpha = 0.05 for a two-tailed test) estimated by 200kb weighted block jackknife. Grey dashed lines represent the PBSnj genic enrichment in simulations, under increasing levels of background selection that best match different aspects of δ, as described in Figs. 2 and 3: lightest to darkest shades: B= 0.925 (excluding δ tail bins), 0.888 (all δ bins), and 0.863 (only δ tail bins). **b,** BGS does not result in eastern and central chimpanzees differing in the PBSnj tail bin genic enrichment. X axis is the log_2_ ratio of the PBSnj tail genic enrichment eastern / pop2. Grey shaded ticks represent the PBSnj genic enrichment in simulations, under increasing levels of background selection that best match different aspects of *δ*, as described in Figs. 2 and 3: lightest to darkest shades: B= 0.925 (excluding *δ* tail bins), 0.888 (all *δ* bins), and 0.863 (only *δ* tail bins).

**Figure 4-figure supplement 3:**
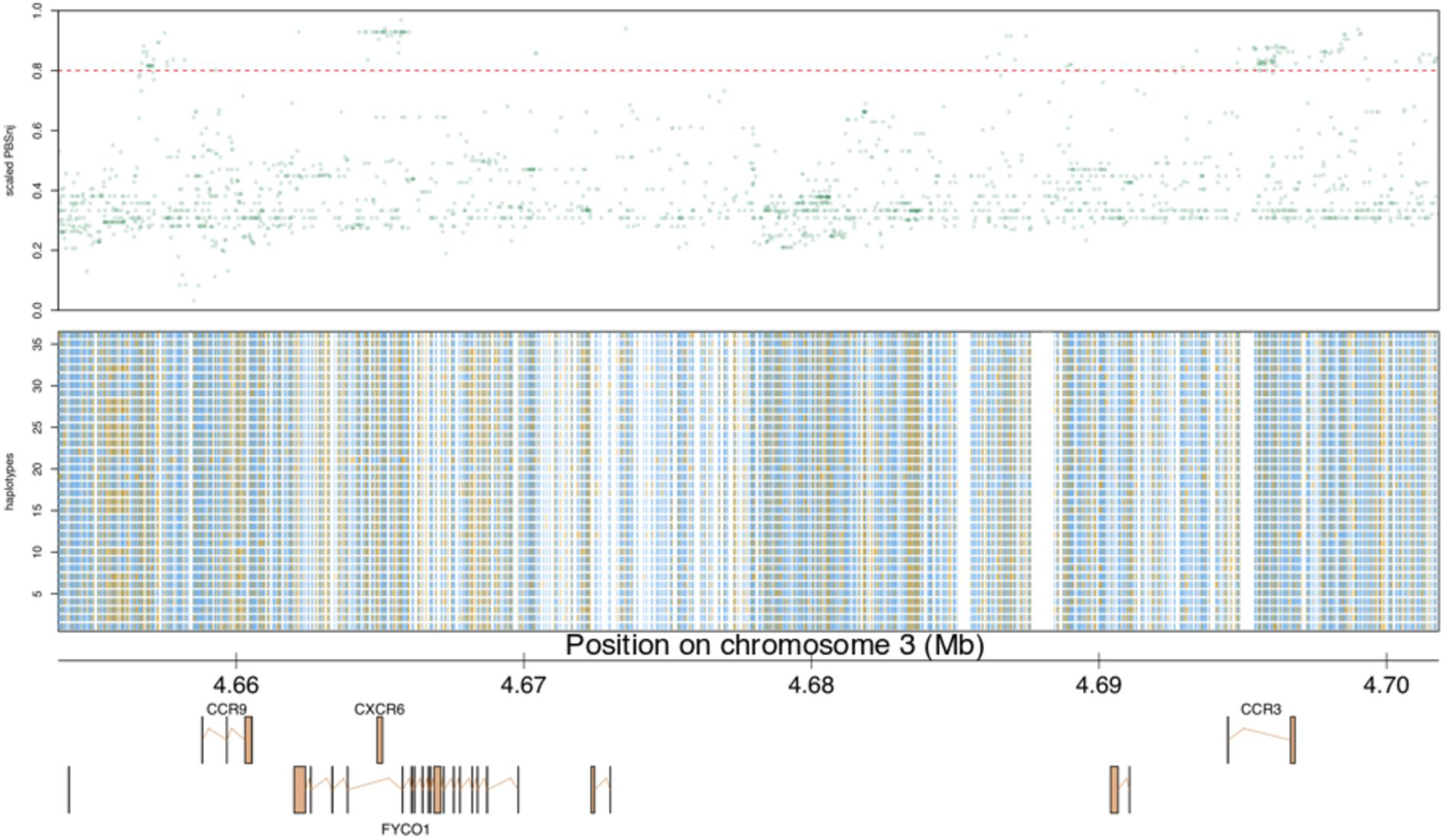
Number of sweeps in the chromosome 3 chemokine receptor cluster of central chimpanzees. X axis: position along chromosome 3 (Mb). Plotted in the upper panel are the PBSnj central scores in the region encompassing *CCR3*, *CCR9*, and *CXCR6*. An independent cluster of high PBSnj scores is associated with each candidate gene. Each point represents one PBSnj score, colour has an alpha = 30% to reduce over plotting. Haplotypes are plotted in the central panel. Yellow ticks are derived alleles, blue are ancestral, while white is space so that each tick aligns with PBSnj scores. Inspection indicates that there is a degree of haplotype scrambling between each of the candidate genes. Lastly, we depict the genes in this region in the lower panel.

**Appendix 1 Figure 1:**
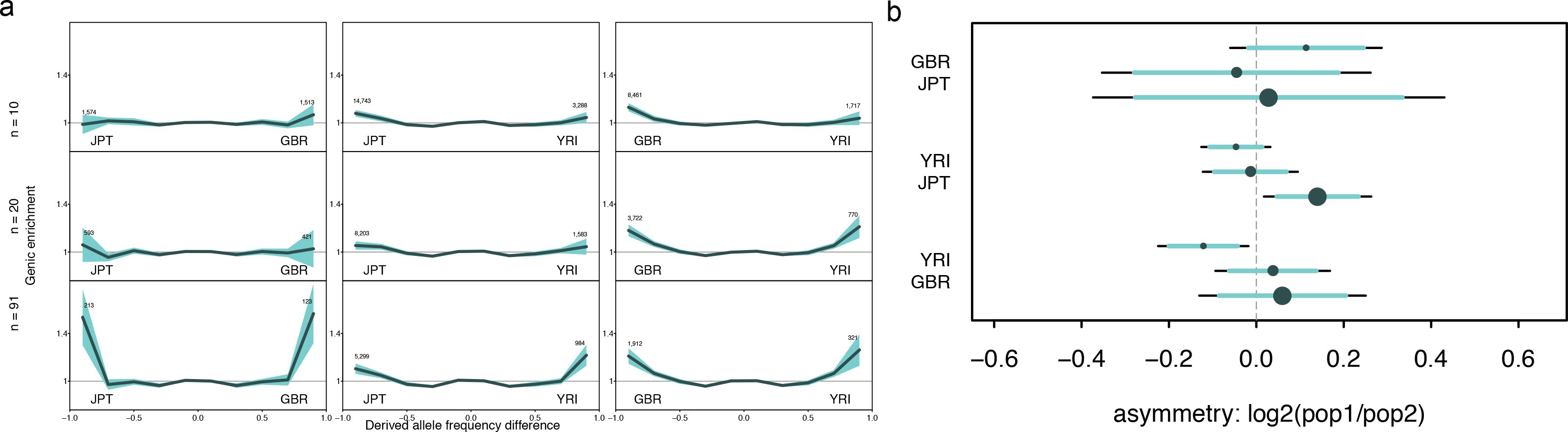
Genic enrichment in bins of signed difference in derived allele frequency (δ), for human populations from the 1000 Genomes Phase III. ***a,*** X-axis: *δ* is computed as the difference in derived allele frequency, for each pair of populations. Tail bins (the last bin in either end of *δ*) contain those SNPs with the largest allele frequency differences. Numbers are of the genic SNPs in each tail bin. Y-axis: genic enrichment in each *δ* bin, computed as described in Methods. Shading represents the 95% CI (i.e. alpha = 0.05 for a two-tailed test) estimated by 200kb weighted block jackknife**, b,** The asymmetry of the genic enrichments in the *δ* tails is measured by taking their log_2_ ratio, thus 0 indicates a symmetric enrichment (equal enrichment in both *δ* tails). Dot = observed asymmetry, with size indicating the relative sample size (10, 20, 91 individuals). Horizontal lines represent confidence intervals estimated by 200kb weighted block jackknife (light = 95%, black = 99%, i.e. alpha = 0.05 or 0.01 for a two-tailed test).

**Appendix 3 Figure 1:**
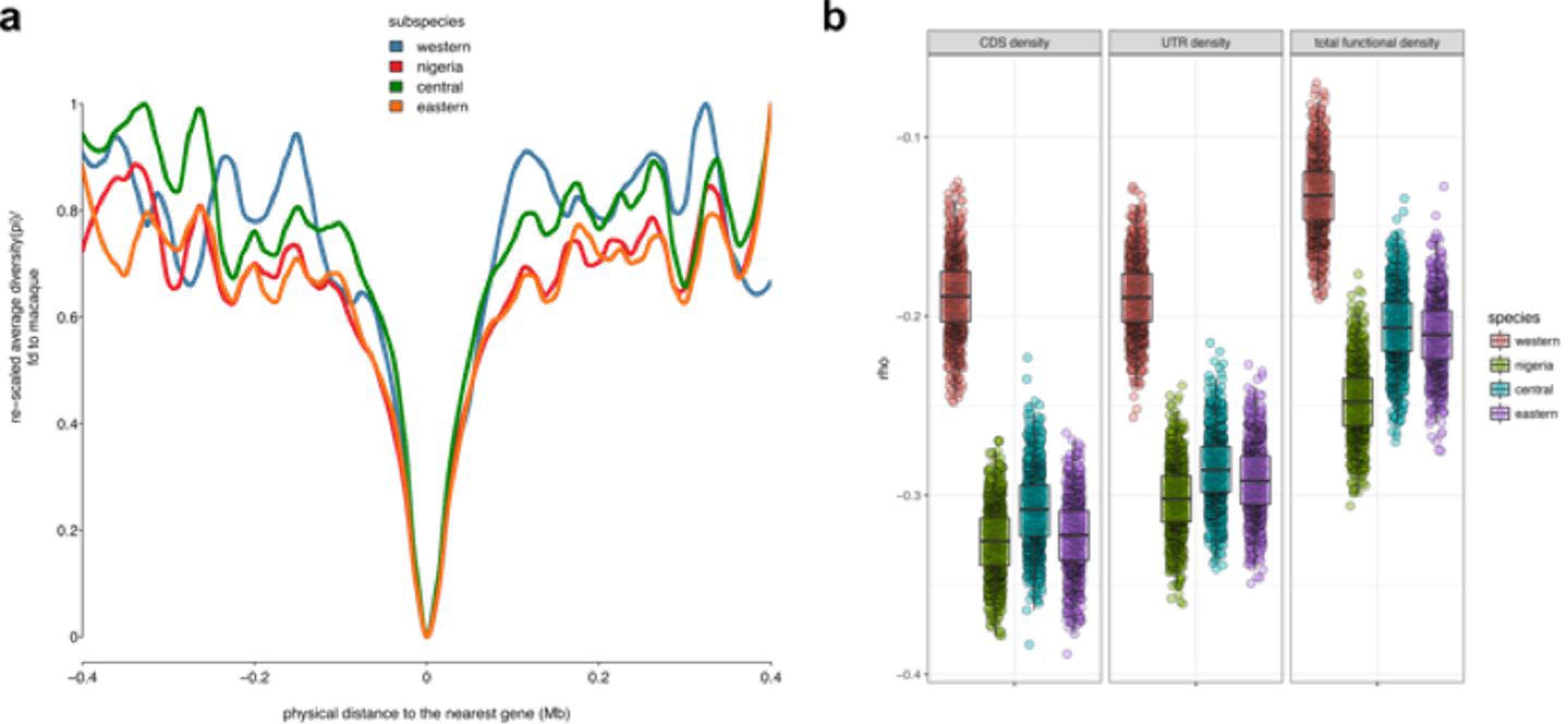
The effect of background selection on patterns of neutral diversity in chimpanzees. **a**, Diversity levels at neutral sites as a function of the distance to the nearest gene. We calculated scale diversity (pi / divergence to macaque) in bins of distance to genic regions. We then rescaled scaled diversity for each subspecies so that the diversity was in the range 0-1. **b**, To further explore the effects of BGS on chimpanzee genomes we checked the correlation of density of functional sites with neutral diversity (pi). We used windows 500kb spaced at least 1MB apart in the genome. Here, *rho* is the spearman rank partial correlation between windowed diversity and density of functional sites per window, controlling for recombination rate (the average rate per window). Each dot represents a bootstrap replicate (random sample of 500 kb windows). We calculated the partial rho for each bootstrap. Box plots show the median and interquartile ranges of the bootstrap replicates.

**Appendix 4 Figure 1:**
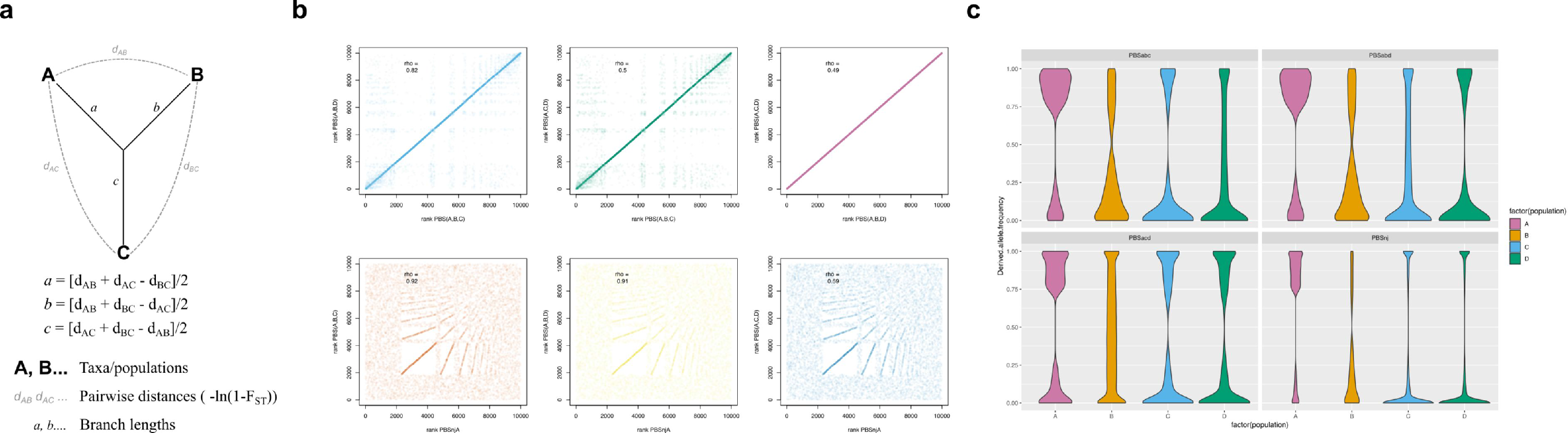
Deriving the PBSnj statistic. **a,** PBS is just a simple arithmetic function of pairwise F_ST_ values for a group of three taxa or populations. **b,** The configuration or choice of populations determines the information content of PBS. In each panel are the spearman’s rho correlations between different PBS configurations, and between PBS and our new statistic PBSnj for a simple four population model (described in Appendix 4). In each case Pop A is the focal population. PBS_ABC_ and PBS_ABD_ are highly correlated but not identical indicating that incorporating both Pops C and D would refine the identification of Pop A specific differentiated variants. PBSnjA, which utilises information from all four populations is more highly correlated with both PBS_ABC_ and PBS_ABD_ than they are with each other. Alpha = 10% for plotted points to reduce over saturation. **c,** For each statistic, we plot the site frequency spectrum (SFS) for each of the four populations for sites identified as outliers in Pop A. PBSnj clearly finds those sites differentiated in Pop A, and better than either PBS_ABC_ and PBS_ABD_. In the standard PBS, the SFS in the species not included in the PBS configuration has a more uniform distribution, indicating that some sites identified as PBS frequency outliers in Pop A are not true population specific outliers.

**Appendix 5 Figure 1:**
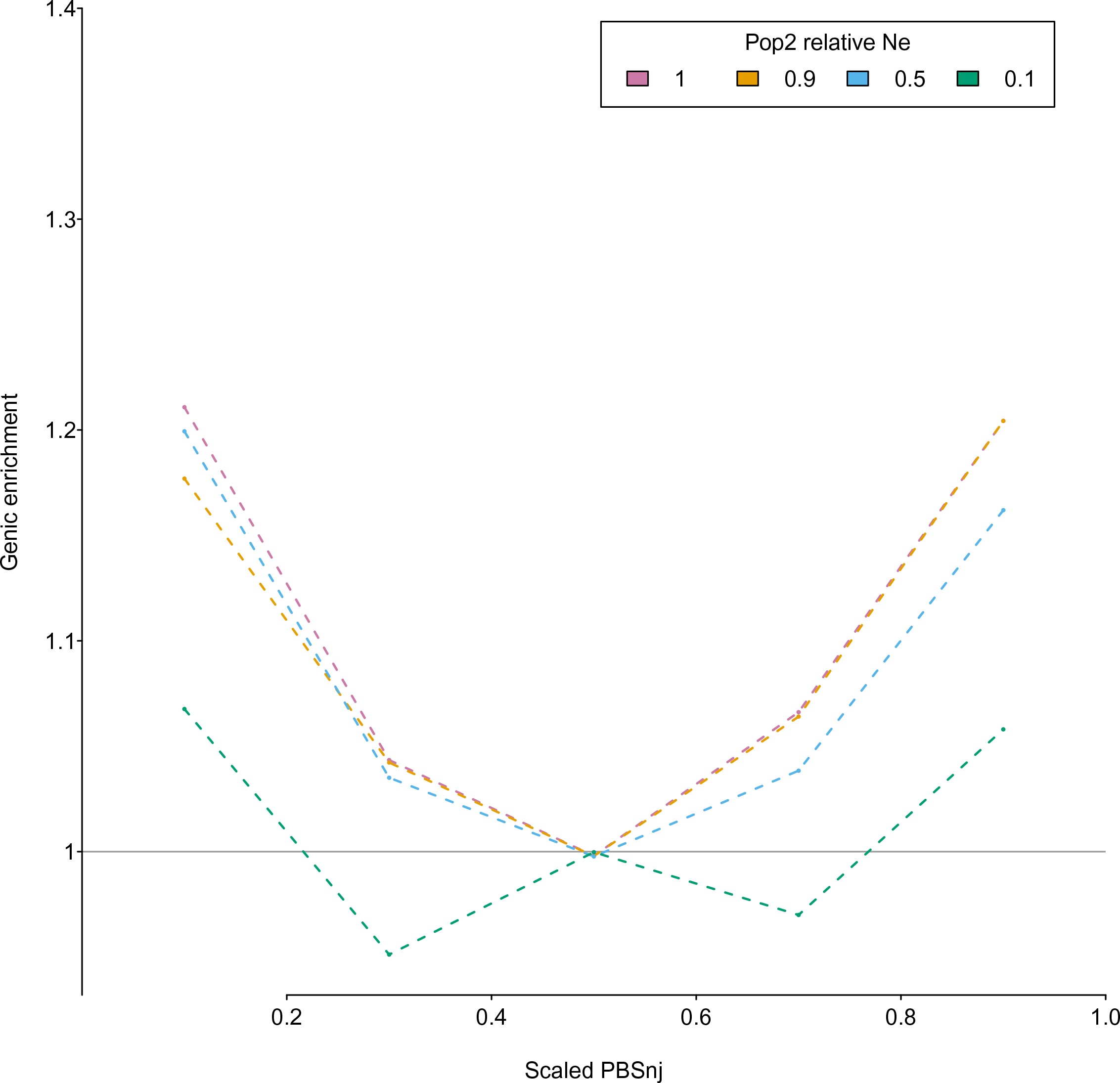
Effect of reduced Ne on PBSnj genic enrichments. In a simple four population model, we modelled genic regions as those with a B = 0.9. In population 2, we simulated four effective population size ratios (1, 0.9, 0.5, 0.1). *N_e_* ratios of 0.5 and 0.1 result in a reduced genic enrichment given the same strength of background selection. X-axes: PBS scaled to take values in the range 0 −1, per subspecies. Y-axes: Genic enrichment.

## Source data files

Figure 1–Source Data 1: Genome wide average heterozygosity counts for chimpanzees.

Figure 2–Source Data 1: Observed and BGS simulated Genic enrichment in bins of signed difference in derived allele frequency.

Figure 3–Source Data 1: Observed and BGS simulated Genic log 2 ratios of tail bin enrichment, signed difference in derived allele frequency.

Figure 3-figure supplement 1–Source Data 1: Observed and BGS simulated, with greater strength of BGS in eastern chimpanzees, Genic log 2 ratios of tail bin enrichment, signed difference in derived allele frequency.

Figure 4–Source Data 1: Genic enrichment in bins of PBSnj in eastern and central chimpanzees

Figure 4-figure supplement 2-Source data 1 Genic enrichment in bins of PBSnj in all four chimpanzee sub-species.

## Supplementary Files

Supplementary File 1: Signed difference in derived allele frequency, genic and non-genic tail counts.

Supplementary File 2: Observed and simulated *δ* bin genic enrichments

Supplementary File 3: Observed and model chimpanzee subspecies F_ST_

Supplementary File 4: Fit of simulated to observed genic enrichments across δ bins.

Supplementary File 5: log_2_ ratio of eastern and central chimpanzee *δ* tail bin genic enrichments with different strengths of background selection.

Supplementary File 6: Model-based reduction of neutral diversity in chimpanzee sub-species. Models are tested for their ability to explain diversity as a function of distance to functional sites.

Supplementary File 7: Effect of divergence on δ tail SNP number.

Supplementary File 8: Effect of *N_e_* on δ tail SNP number.

Supplementary File 9: Fitting BGS to match observed PBSnj tail genic enrichments.

Supplementary File 10: PBSnj tail SNP haplotype statistic scores.

Supplementary File 11: Non-synonymous PBSnj eastern tail SNPs.

Supplementary File 12: PBSnj tail SNP regulomeDB enrichments.

Supplementary File 13: PBSnj tail SNP conservation/phastCons score enrichments.

Supplementary File 14: Effect of Ancestral Allele estimation on eastern vs. central chimpanzee δ bin genic enrichments.

Supplementary File 15: PBSnj Eastern GO enrichment.

Supplementary File 16: PBSnj Central GO enrichment.

Supplementary File 17: PBSnj Nigeria-Cameroon GO enrichment.

Supplementary File 18: PBSnj Western GO enrichment.

Supplementary file 19: PBSnj Eastern VIP enrichment.

Supplementary file 20: PBSnj Central VIP enrichment.

Supplementary file 21: PBSnj Nigeria-Cameroon VIP enrichment.

Supplementary file 22: PBSnj Western VIP enrichment.

Supplementary file 23: SIV responsive gene enrichment tests.

Supplementary File 24: PBSnj eastern tail and SIV responsive genes SNP regulomeDB enrichment

Supplementary file 25: PBSnj Eastern and SIV gene GO enrichment

## Source code

### PBSnj_method.R

Contains function for calculating PBSnj from the full pairwise F_ST_ matrix of four populations.

